# A Guide to Pre-Processing High-Throughput Animal Tracking Data

**DOI:** 10.1101/2020.12.15.422876

**Authors:** Pratik Rajan Gupte, Christine E. Beardsworth, Orr Spiegel, Emmanuel Lourie, Sivan Toledo, Ran Nathan, Allert I. Bijleveld

## Abstract

1. Modern, high-throughput animal tracking studies collect increasingly large volumes of data at very fine temporal scales. At these scales, location error can exceed the animal’s step size, leading to mis-estimation of key movement metrics such as speed. ‘Cleaning’ the data to reduce location errors prior to analyses is one of the main ways movement ecologists deal with noisy data, and has the advantage of being more scalable to massive datasets than more complex methods. Though data cleaning is widely recommended, and ecologists routinely consider cleaned data to be the ground-truth, inclusive uniform guidance on this crucial step, and on how to organise the cleaning of massive datasets, is still rather scarce.
2. A pipeline for cleaning massive high-throughput datasets must balance ease of use and computationally efficient signal vs. noise screening, in which location errors are rejected without discarding valid animal movements. Another useful feature of a pre-processing pipeline is efficiently segmenting and clustering location data for statistical methods, while also being scalable to large datasets and robust to imperfect sampling. Manual methods being prohibitively time consuming, and to boost reproducibility, a robust pre-processing pipeline must be automated.
3. In this article we provide guidance on building pipelines for pre-processing high-throughput animal tracking data in order to prepare it for subsequent analysis. Our recommended pipeline, consisting of removing outliers, smoothing the filtered result, and thinning it to a uniform sampling interval, is applicable to many massive tracking datasets. We apply this pipeline to simulated movement data with location errors, and also show a case study of how large volumes of cleaned data can be transformed into biologically meaningful ‘residence patches’, for quick biological inference on animal space use. We use calibration data to illustrate how pre-processing improves its quality, and to verify that the residence patch synthesis accurately captures animal space use. Finally, turning to tracking data from Egyptian fruit bats (*Rousettus aegyptiacus*), we demonstrate the pre-processing pipeline and residence patch method in a fully worked out example.
4. To help with fast implementation of standardised methods, we developed the `R` package `atlastools`, which we also introduce here. Our pre-processing pipeline and `atlastools` can be used with any high-throughput animal movement data in which the high data-volume combined with knowledge of the tracked individuals’ movement capacity can be used to reduce location errors. The `atlastools` function is easy to use for beginners, while providing a template for further development. The use of common pre-processing steps that are simple yet robust promotes standardised methods in the field of movement ecology and leads to better inferences from data.

## 1 Introduction

The movement of an animal is an adaptive, integrated response to multiple drivers, including internal state, life-history traits and capacities, biotic interactions, and other environmental factors (Holyoak et al., 2008; Nathan et al., 2008). The movement ecology framework links the drivers, processes, and fitness outcomes of animal movement (Nathan et al., 2008), and remotely tracking individual animals in the wild is the methodological mainstay of movement ecology (Hussey et al., 2015; Kays et al., 2015; Nathan et al., 2008; Wikelski et al., 2007). A key challenge with observed tracks is to extract information on the behavioural, cognitive, social, ecological and evolutionary processes that shape animal movement. Tracking data, which are observations of a continuous process (animal movement) at discrete timesteps, reveal useful information about the movement process when the tracking interval is considerably shorter than the typical duration of a movement mode (Getz and Saltz, 2008; Nathan et al., 2008; Noonan et al., 2019). This can be accomplished by wildlife tracking systems that collect position data from many individuals at high temporal and spatial resolution (i.e., high-throughput tracking) relative to the scale of the movement mode of interest (Getz and Saltz, 2008). High-throughput tracking technologies include GPS tags (Harel et al., 2016; Klarevas-Irby et al., 2021; Papageorgiou et al., 2019; Strandburg-Peshkin et al., 2015), tracking radars (Horvitz et al., 2014), and computer vision methods for tracking entire groups of animals from video recordings (Pérez-Escudero et al., 2014; Rathore et al., 2020). Furthermore, high-throughput wildlife tracking is routinely provided by ‘reverse-GPS’ systems developed to track animals over land, such as ATLAS (Advanced Tracking and Localization of Animals in real-life Systems Toledo et al., 2014, 2016, 2020; Weiser et al., 2016, see also MacCurdy et al. 2009, 2019), and acoustic tracking of aquatic animals (Aspillaga et al., 2021a,b; Baktoft et al., 2019, 2017; Jung et al., 2015). Finally, low resolution tracking over a long duration may also capture important aspects of animal behaviour at certain time-scales (e.g. migration, long-range dispersal; Getz and Saltz, 2008), thereby being ‘relatively’ high-throughput.

Although high-throughput tracking provides a massive amount of data on the path of a tracked animal, these data present a challenge to ecologists. When tracking animals at a high temporal resolution, the location error of each position may approach or exceed the true movement distance of the animal, compared to low-resolution tracking with the same measurement error. This leads to an over-estimation of the true distance moved by an animal between two discrete time-points, leading to unreliable metrics ultimately derived from movement distance, such as speed and tortuosity (see Calenge et al., 2009; Hurford, 2009; Noonan et al., 2019; Ranacher et al., 2016). Additionally, the location error around a position introduces uncertainty when studying the relationship between animal movements and either fixed landscape features (e.g. roads) or mobile elements (e.g. other tracked individuals). Users have two main options to improve data quality, *(1)* making inferences after modelling the system-specific location error using a continuous time movement model (Aspillaga et al., 2021<u>b</u>; Fleming et al., 2014, 2020; Johnson et al., 2008; Jonsen et al., 2005, 2003; Patterson et al., 2008), or *(2)* pre-processing data to clean it of positions with large location errors (Bjørneraas et al., 2010). The first approach may be limited by the animal movement models that can be fitted to the data (Fleming et al., 2014, 2020; Noonan et al., 2019), may result in unreasonable computation times, or may be entirely beyond the computational capacity of common hardware, leading users to prefer data cleaning instead. Data cleaning reveals another challenge of high-throughput tracking: the large number of observations make it difficult for researchers to visually examine each animal’s track for errors (Toledo et al., 2020; Weiser et al., 2016). With manual identification and removal of errors from individual tracks prohibitively time consuming, data cleaning can benefit from automation based on a protocol.

Pre-processing of movement data — defined as the set of data management steps executed prior to data analysis — must reliably discard large location errors, also called outliers, from tracks (analogous to reducing false positives) while avoiding the overzealous rejection of valid animal movements (analogous to reducing false negatives). How well researchers balance these imperatives has consequences for downstream analyses (Stine and Hunsaker, 2001). For instance, small-scale resource selection functions can easily infer spurious preference and avoidance effects when there is uncertainty about an animal’s true position (Visscher, 2006). Ecologists recognise that tracking data are imperfect observations of the underlying movement process, yet they implicitly consider cleaned data equivalent to the ground-truth. This assumption is reflected in popular statistical methods in movement ecology such as Hidden Markov Models (HMMs) (Langrock et al., 2012), stationary-phase identification methods (Patin et al., 2020), or step-selection functions (SSFs) (Avgar et al., 2016; Barnett and Moorcroft, 2008; Signer et al., 2017), which expect minimal location errors relative to real animal movement (i.e., a high signal-to-noise ratio). This makes the reproducible, standardised removal of location errors crucial to any animal tracking study. While gross errors are often removed by positioning-system algorithms, ‘reasonable’ errors often remain to confront end users (Fischler and Bolles, 1981; Ranacher et al., 2016; Weiser et al., 2016). Further, as high-throughput tracking is deployed in more regions and for more species, standardised pre-processing steps should be general enough to tackle animal movement data recovered from a range of environments, so as to enable sound comparisons across species and ecosystems.

Despite the importance and ubiquity of reducing location errors in tracking data, movement ecologists lack formal guidance on this crucial step. Pre-processing protocols are not often reported in the literature, or may not be easily tractable for mainstream computing hardware and software. Some tracking data, such as GPS, are autonomously pre-processed without user access to the raw data (using error estimates and Kalman smooths; Kaplan and Hegarty, 2005, and substantial location errors may yet persist). However, filtering out positions using estimates of location error alone may not be sufficient to exclude outliers which represent unrealistic movement but have low error measures (Ranacher et al., 2016; Weiser et al., 2016). When tracking systems do make their raw data available to researchers, this can enable users to better control the data pre-processing stage, and to substantially improve data quality while ensuring that cleaning does not itself lead to unrealistic movement tracks (e.g. Kalman smooths which distort tracks, Kaplan and Hegarty 2005). Furthermore, This makes identifying and removing biologically implausible locations from a track an important component of recovering true animal movement (Bjørneraas et al., 2010). Even after removing unrealistic movement, a track may be comprised of positions that are randomly distributed around the true animal location (Noonan et al., 2019). The large data-volumes of high-throughput tracking allow for a neat solution: tracks can be ‘median smoothed’ to reduce small location errors that have remained undetected. Large data volumes may also need to be thinned, for example, examining environmental covariates as predictors of prolonged residence in an area (see e.g. Aarts et al., 2008; Bijleveld et al., 2016; Bracis et al., 2018; Harel et al., 2016; Oudman et al., 2018) might require thinning of high-resolution movement data to match the lower spatial resolution of environmental measurements. Data thinning and clustering are also required to avoid non-independent observations due to strong spatio-temporal autocorrelation, or to examine the effect of sampling scale on movement metrics and resource-selection (Fleming et al., 2014; Noonan et al., 2019).

When dealing with datasets that contain many millions of positions, reseachers may run into computational limits when trying to apply pre-processing steps to their full dataset. For instance, the size of working memory (RAM) limits the size of datasets that can be loaded into R, the programming and statistical language of choice in movement ecology (Joo et al., 2020a,b; R Core Team, 2020). Data-rich fields such as genomics inspire a possible solution: to break very large data into smaller subsets, and pass these subsets through automated computational ‘pipelines’ (Peng, 2011; Schadt et al., 2010). Building a pre-processing pipeline for movement data may be expected to comprise of three primary concerns: *(1)* identifying necessary pre-processing steps, *(2)* quality-control on the steps and the resultant data, and *(3)* increasing the speed and efficiency of the pipeline. While exploratory data analysis and visualisation can help determine how to pre-process the data to maximise the signal to noise ratio (Slingsby and van Loon, 2016), standardising implementations of pre-processing techniques into robust, version controlled software packages (e.g. in R, see Wickham, 2015), is key to increasing the reliability and reproducibility of animal movement ecology (Archmiller et al., 2020; Haddaway and Verhoeven, 2015; Lewis et al., 2018; Powers and Hampton, 2019). Overcoming hard computational constraints on speed and memory usage for very large data will often require a combination of programming strategies, such as using tools optimised for tabular data, or parallelised processing.

Here, we present guidelines for building reproducible pre-processing pipelines for high-throughput tracking data (Fig. 1), with a focus on simple, widely generalisable pre-processing steps that help improve data quality (Fig. 2). We take two important considerations into account, that *(1)* the pre-processing steps should be easily understood and reproduced, and *(2)* our implementations must be computationally efficient and reliable. Consequently, formalising tools as functions in an `R` package would improve portability and reproducibility (Marwick et al., 2018; Wickham, 2015). We demonstrate simple yet robust implementations of the pre-processing steps we recommend, conveniently wrapped into the `R` package `atlastools` (Gupte, 2020), with a discussion of features that make these steps more reproducible, and more efficient. In two fully worked out examples using our package on tracking data, we show how to apply basic spatio-temporal and data quality filters, how to filter out unrealistic movement, and how to reduce the effect of location error with a median smooth. Then, we suggest one potential application of high-throughput tracking in studies of animal movement and space use, illustrated by the first-principles based synthesis of ‘residence patches’ from clusters of spatio-temporally proximate positions (*sensu* Barraquand and Benhamou, 2008; Bijleveld et al., 2016; Oudman et al., 2018). Using calibration data from an ATLAS system, we show how the residence patch segmentation-clustering method can be used to accurately identify areas of prolonged residence under real field conditions. Finally, we use ATLAS data from Egyptian fruit bats (*Rousettus aegyptiacus*) tracked in the Hula Valley, Israel, to show a fully worked out example of the pre-processing pipeline and the residence patch method.

**Figure 1.**
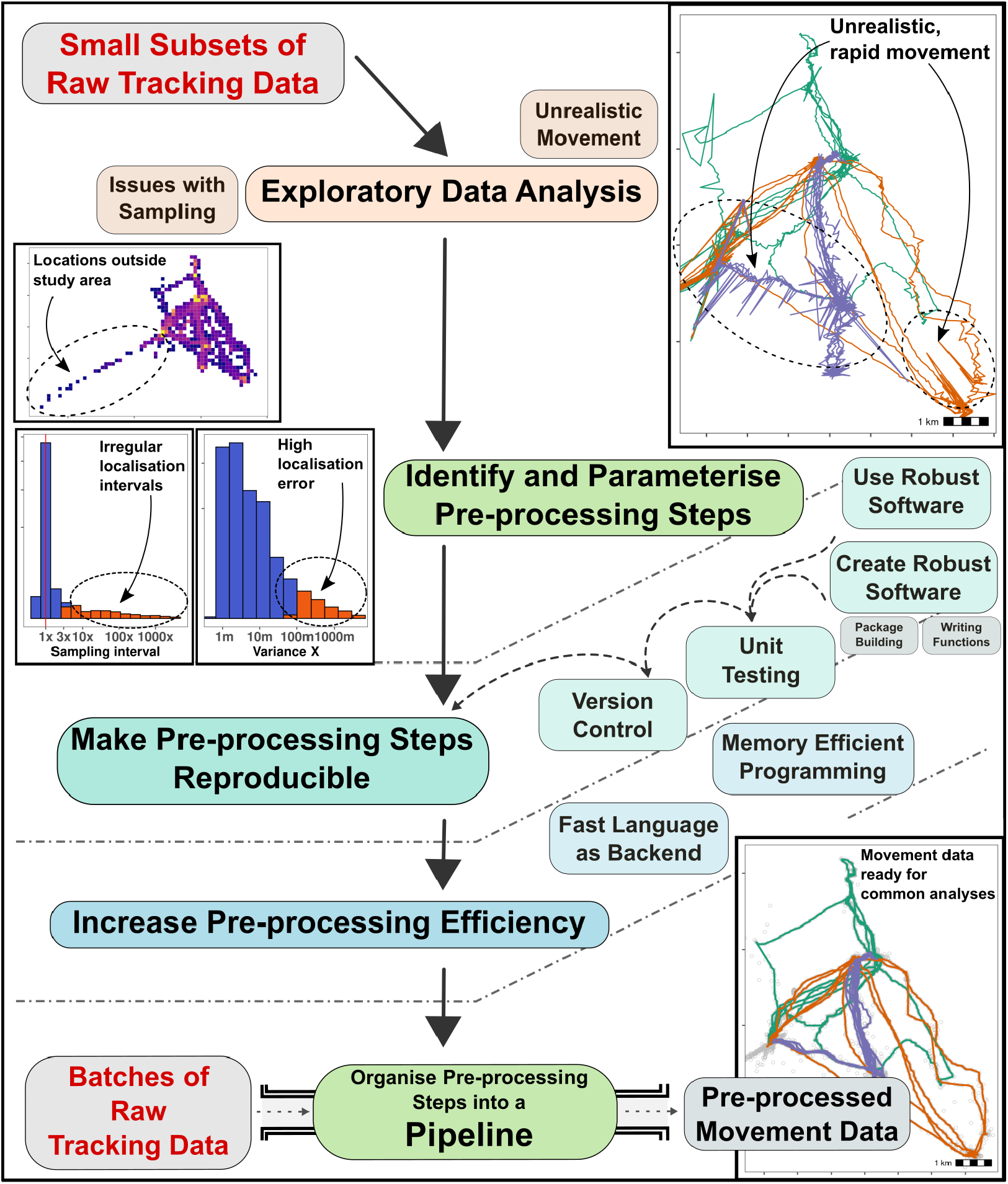
A general workflow for approaching the pre-processing of very large, high-throughput tracking datasets. Researchers should begin with exploratory data analysis of representative subsets of the data to identify common issues with quality, and use EDA to select and parameterise pre-processing steps such as filters and smooths (*see Fig. 2*). Pre-processing steps implemented as programming code can be made reproducible and shareable by following best-practices for software development: (1) encapsulating code as functions, (2) bundling related functions into a package, (3) testing the functions to ensure they work as expected, and (4) implementing version control, using a system such as git to clarify which version was used in an analysis. The efficiency of pre-processing tools can be increased by using existing tools optimised for large datasets, or by writing new tools (code) in a ‘fast’ language such as C++ (see main text for examples). Finally, batches of the full dataset can be passed through a pipeline comprised of the pre-processing tools to obtain cleaned movement data, suitable for a range of analyses. Fig. 2 shows an example of such a pipeline.

**Figure 2.**
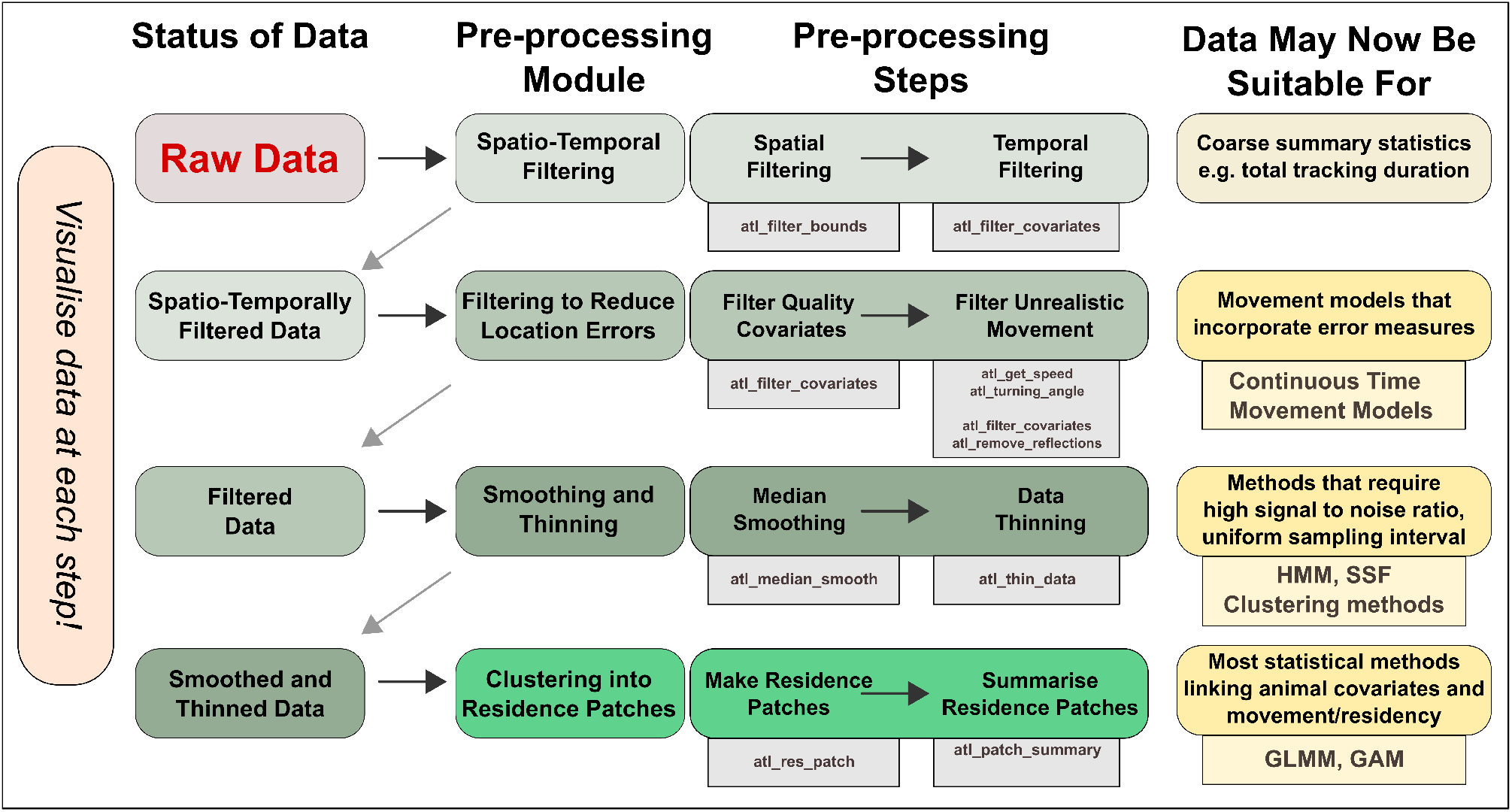
A general, modular pipeline for pre-processing high-throughput tracking data from raw localisations to cleaned data, and optionally into residence patches. Users should apply the appropriate pre-processing modules and the steps therein until the data are suitable for their intended analysis, some of which are suggested here. The `atlastools` function that may be used to implement each pre-processing step is shown in the grey boxes underneath each step. Popular statistical methods are shown underneath possible analyses (yellow boxes). Users are strongly encouraged to visualise their data and scan it for location errors as they work through the pipeline, always asking the question, could the animal plausibly move this way?

## 2 Building Pre-Processing Pipelines for Large Tracking Data

### 2.1 Exploratory Data Analysis to Identify Pre-processing Steps

Exploratory data analysis (EDA) is the first step towards building a data pre-processing pipeline (see Fig. 1; Slingsby and van Loon, 2016). Researchers with very large datasets of perhaps millions of rows should ideally select a representative subset of these data for EDA, including individuals of different species, sexes, or seasonal cohorts. Examples of EDA include plotting heatmaps of the number of observations per unit area across the study site (Fig. 1). Histograms of the the location error estimates, plotting the linear approximations of animal paths between observations, and histograms of the sampling interval can help determine how data need to be treated so as to minimise location errors and improve computational tractability (Fig. 1). While pre-processing steps required for datasets will differ between studies and tracking technologies, we elaborate upon candidate steps and their parameterisation in following sections (see also Fig. 2).

### 2.2 Improving Reliability and Reproducibility

Following EDA and the parameterisation of data cleaning steps, researchers must prioritise making their implementation of these steps reliable and reproducible (Fig. 1; see also Alston and Rick, 2021). Reproducing pre-processing steps can be challenging when using only written descriptions from published articles. Providing the code to implement pre-processing steps reduces ambiguity and increases reproducibility (Haddaway and Verhoeven, 2015). The best-practices here are *(1)* to implement pre-processing steps as ‘functions’, *(2)* to collect related functions — e.g. for similar kinds of data — into a software ‘package’, *(3)* to ‘test’ that the functions handle input as expected, and *(4)* implement ‘version control’ throughout, such that the process of development is documented (Fig. 1; Alston and Rick, 2021; Perez-Riverol et al., 2016; Wickham, 2015). The `atlastools` package incorporates these best-practices, and may be used as a reference (Gupte, 2020). We have written each pre-processing step as a separate function, and each of these functions is tested, usually on simulated data, but in some cases also on empirical data (Wickham, 2015, see the directory `tests/` in the associated Zenodo repository). Other good examples of `R` packages following these best-practices are ‘move’ (Kranstauber et al., 2011), and ‘sftrack’ (Boone et al., 2020).

### 2.3 Improving Speed and Efficiency

Animal tracking data stored in a relational database (e.g. SQL databases Codd, 1970) naturally lends itself to batch-processing, as the data can be broken into meaningful subsets based on individual identity and tracking season. These smaller subsets can then be loaded into working memory, pre-processed, and saved in a separate location (see Supplementary Material 1, Section 2 for a worked out example on an SQL database). Batch-processing allows a rudimentary form of parallel-processing, by allowing each subset of the data to be processed independently of the full dataset, for example, using a computing cluster (see also Dai, 2021, for an alternative). While cluster-computing and other advanced methods can lead to significant speed gains, they may be challenging to implement for beginning practitioners. Using existing tools optimised for tabular data, such as the `data.table` `R` package (Dowle and Srinivasan, 2020), can also speed up computation; `atlastools` is built using `data.table` for this reason. Finally, a relatively advanced method used by packages such as `move` and `recurse` (Bracis et al., 2018) is to write one’s own methods in a ‘fast’ low-level language, such as `C++`, and link these to `R` (Eddelbuettel, 2013).

## 3 Pre-processing Steps, Usage, and Simulating Data

### 3.1 Pre-processing Steps and atlastools

We lay out pre-processing techniques for raw high-throughput tracking data, and demonstrate working examples of these techniques, which we have collected in the `R` package `atlastools` (Fig. 2). Our package is aimed getting ‘raw data’ to the ‘analysis’ stage identified by Joo et al. (2020) in their review of R packages in movement ecology. The package is based on `data.table`, a fast implementation of data frames; thus it is compatible with a number of data structures from popular packages including `move`, `sftrack`, and `ltraj` objects, which can be converted to data frames (Boone et al., 2020; Calenge et al., 2009; Kranstauber et al., 2011). These pre-processing techniques and package were designed with ATLAS systems in mind, motivated to meet the rapid growth of studies using this high-throughput system worldwide: in Israel (Corl et al., 2020; Toledo et al., 2014, 2016, 2020; Vilk et al., 2021), the UK (Beardsworth et al., 2021b,c), and the Netherlands (Beardsworth et al., 2021a). However, the principles and functions presented here are ready for use with other massive high-resolution data collected by GPS, reverse-GPS or any other high-throughput tracking system. Users may construct a pre-processing pipeline comprising of all the techniques we cover, or implement the modules most suitable for their data. Users are advised to visualise their data throughout their workflow, and especially to perform thorough EDA, to check for evident location errors or other issues (Slingsby and van Loon, 2016).

1. Users can use simple spatio-temporal filters to select positions within a certain time or area.
2. Next, users should reduce gross location errors by removing unreliable positions based on a system-specific error measure, or by the plausibility of associated movement metrics, such as speed and turning-angle (Calenge et al., 2009; Seidel et al., 2018).
3. Users should then reduce small-scale location errors by applying a median smooth.
4. Users who need uniformly thinned data can either aggregate or subsample it. At this stage, the data are ready for a number of popular statistical treatments such as Hidden Markov Model-based classification (Langrock et al., 2012; Michelot et al., 2016).
5. Finally, users wishing simple, efficient segmentation-clustering of points where the animal showed prolonged residence, can classify their data into ‘residence patches’ (Barraquand and Benhamou, 2008; Bijleveld et al., 2016) based on the movement ecology of their study species, after filtering out travelling segments (see Segmenting and Clustering Movement Tracks into Residence Patches).

### 3.2 Demonstrating Pre-processing Steps with Simulated Data

To demonstrate pre-processing steps, we simulated a realistic movement track of 5,000 positions using an unbiased correlated velocity model (UCVM) implemented via the R package smoove (Gurarie et al., 2017, see Fig. 2.a). We added three kinds of error to the simulated track: (1) normally distributed small-scale offsets to the X and Y coordinates independently, (2) normally distributed large-scale offsets to a random subset (0.5 %) of the positions, and (3) large-scale displacement of a continuous sequence of 300 of the 5,000 positions (indices 500 – 800) (Fig. 2.a). To demonstrate the residence patch method, we chose to simulate three independent rotational-advective correlated velocity movement (RACVM) tracks of 500 positions each (*ω* = 7, initial velocity = 0.1, *μ* = 0; see Gurarie et al. 2017), and connected them together with a roughly linear path (see Fig. 5.a). RACVM models can approximate the tracks of soaring birds which circle on thermals over a relatively small area, and move between thermals (‘thermalling’; Gurarie et al., 2017; Harel et al., 2016; Harel and Nathan, 2018). This complex track structure provides a suitable challenge for the residence patch method and helps to demonstrate its generality.

**Listing 1.**
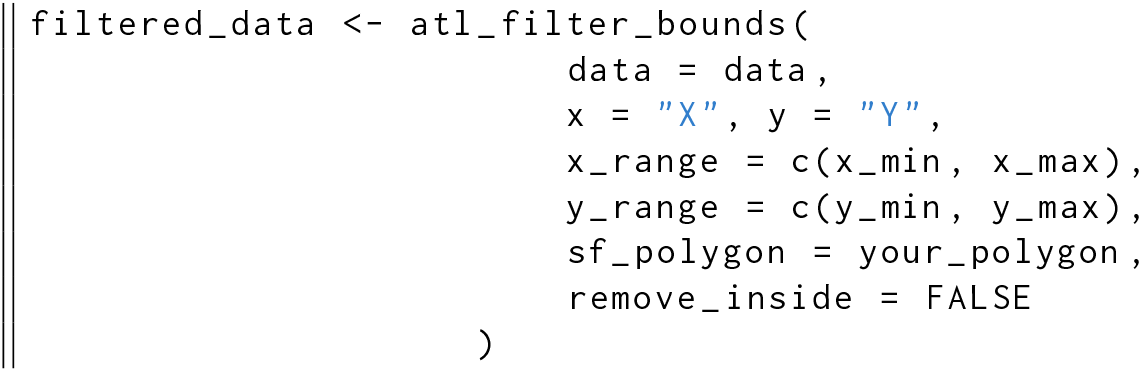
The `atl_filter`_bounds function filters on an area defined by coordinate ranges, a polygon, or all three; it can remove positions outside (`remove_inside = FALSE`), or within the area (`remove_inside = TRUE`). The arguments `x` and `y` determine the X and Y coordinate columns, `x_range` and `y_range` are the filter bounds in a coordinate reference system in metres, and the data can be filtered by an `sf-(MULTI)POLYGON` can be passed using the `sf_polygon` argument. The output is a `data.table`, which must be saved as an object (here, `filtered_data`).

## 4 Spatio-Temporal Filtering

### 4.1 Spatial Filtering Using Bounding Boxes and Polygons

First, users should exclude positions outside the spatial bounds of a study area by comparing position coordinates with the range of acceptable coordinates (the bounding box), and removing positions outside them (Fig. 3.a; Listing 1). A bounding box filter does not require a geospatial representation such as a shapefile, and can help remove unreliable data from a tracking system that is less accurate beyond a certain range (Beardsworth et al., 2021a). In some special cases, users may wish to remove positions *inside* a bounding box, either because movement behaviour within an area is not the focus of a study, or because positions recorded within an area are known to be erroneous. An example of the former is studies of transit behaviour between features which can be approximated by their bounding boxes. Instances of the latter are likely to be system specific, but are known from ATLAS systems. Bounding boxes are typically rectangular, and users seeking to filter for other geometries, such as a circular or irregularly-shaped study area, need a geometric intersection between their data and a spatial representation of the area of interest (e.g. shapefile, geopackage, or `sf`-object in `R`). The `atlastools` function `atl_filter_bounds` implements both bounding box and explicit spatial filters, and accepts X and Y coordinate ranges, an `sf`-polygon or multi-polygon object (Pebesma, 2018), or any combination of the three to filter the data (Listing 1).

**Figure 3.**
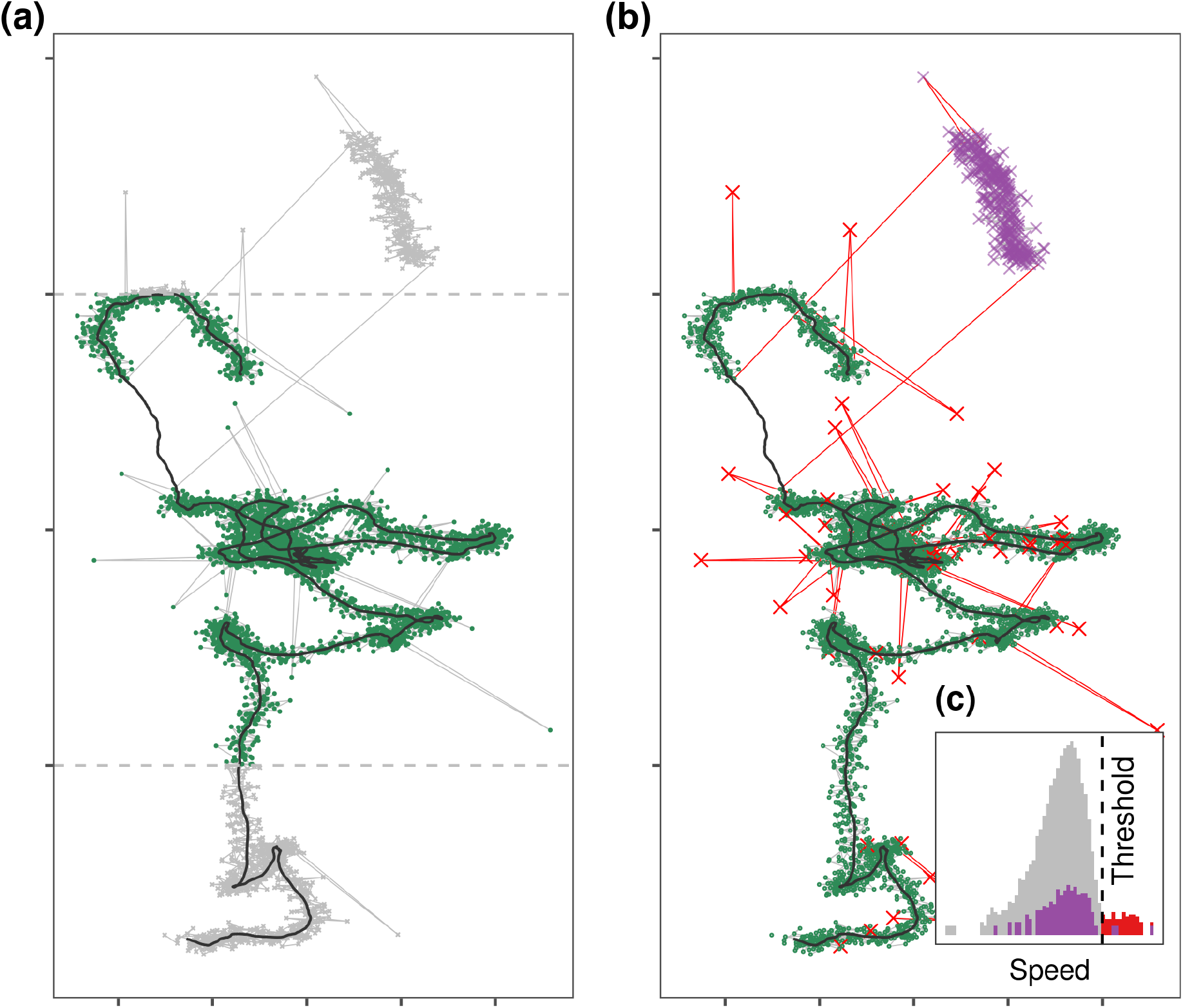
Simulated movement data showing three kinds of artificially added errors. (1) Normally distributed small-scale error on each position, (2) large-scale error added to 0.5% of positions, and (3) 300 consecutive positions displaced to simulate a gross distortion affecting a continuous subset of the track. **(a)** Tracks can be quickly filtered by spatial bounds (dashed grey lines) to exclude broad regions (green = retained; grey = removed). **(b)** location error may affect single observations resulting in point outliers or ‘spikes’ (red crosses and track segments), or continuous subsets of a track, called a ‘prolonged spike’ (purple crosses, top right), and both represent unrealistic movement. **(c)** Histograms of speed for the track (grey = small-scale errors, red = spikes), and the prolonged spike (purple). While spikes can be removed by filtering out positions with high incoming and outgoing speeds and turning angle, prolonged spikes cannot be removed in this way, and should be resolved by conceptualising algorithms that find the bounds of the distortion instead. Users should frequently check the outputs of such algorithms to avoid rejecting valid data.

### 4.2 Temporal and Spatio-temporal Filters

Tracking data might fail to properly represent an animal’s movement at certain times, for instance, data recorded before release, or data from shortly after release when the animal is no influenced by the stress of capture and handling. Periods of poor tracking quality may result from system malfunctions and unusual disturbances, and users may wish to exclude these data as well. Temporal filtering can exclude positions from intervals when data are expected to be unreliable for ecological inference, either due to abnormal movement behaviour or system-specific issues. Temporal filters can be combined with spatial filters to select specific time-location combinations. For example, studies of foraging behaviour of a nocturnal animal would typically exclude tracking data from the animal’s daytime roosts (see Worked out Example, and Supplementary Material 1 Section 2). Users should apply filters in sequence rather than all at once, and visualise the output after each filtering step (‘sanity checks’). The atlastools function `atl_filter_covariates` allows convenient filtering of a dataset by any number of logical statements, including querying data within a spatio-temporal range (Listing 2). The function keeps only those data which satisfy each of the filter conditions, and users must ensure that the filtering variables exist in their dataset in order to avoid errors.

**Listing 2.**
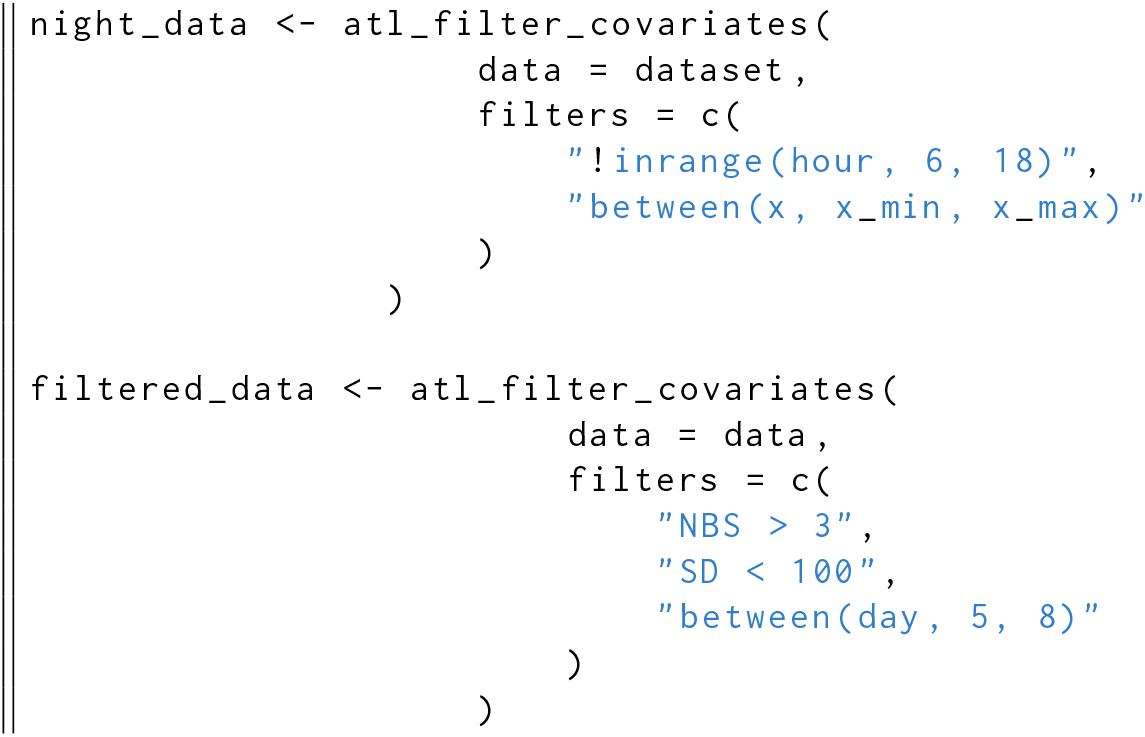
Data can be filtered by a temporal or a spatio-temporal range using `atl_filter_covariates`. Filter conditions are passed to the `filters` argument as a character vector. Only rows in the data satisfying *all* the conditions are retained. Here, the first example shows how nighttime data can be retained using a predicate that determines whether the value of ‘hour’ is between 6 and 18, and also within a range of X coordinates. The second example retains ATLAS locations calculated using > 3 base stations (NBS), with location error (SD) < 100, and data between an arbitrary day 5 and day 8.

## 5 Filtering to Reduce Location Errors

### 5.1 Filtering on Data Quality Attributes

Tracking data attributes can be good indicators of the reliability of positions calculated by a tracking system (Beardsworth et al., 2021a). GPS systems provide direct measures of location error during localisation (Ranacher et al., 2016, Horizontal Dilution of Precision, HDOP in GPS), while reverse-GPS systems provide a similar estimate (called Standard Deviation, SD; MacCurdy et al., 2009, 2019; Ranacher et al., 2016; Weiser et al., 2016). Tracking data can also include indirect indicators of data quality. For instance, GPS systems’ location error may be indicated indirectly by the number of satellites involved in the localisation. In reverse-GPS systems too, the number of base stations involved in each localisation is an indirect indicator of data quality, and positions localised using more receivers are usually more reliable (the minimum required for an ATLAS localisation is 3; see Weiser et al., 2016). Unreliable positions can be removed by filtering on direct or indirect measures of quality using `atl_filter_covariates` (Listing 2).

### 5.2 Filtering Unrealistic Movement

Filtering on system-generated attributes may not remove all erroneous positions, and the remaining data may still include biologically implausible movement. Users are encouraged to visualise their tracks before and after filtering point locations, and especially to ‘join the dots’ and connect consecutive positions with lines (Fig. 3.b). Whether the resulting track looks realistic is ultimately a subjective human judgement, but any decision to filter-out data must remain independent of the hypothesised movement behavior. This basic principle does not preclude explicitly integrating prior knowledge of the movement ecology of the study species to ask, ‘Does the animal move this way?’. Segments which appear to represent unrealistic animal movement are often obvious to researchers with extensive experience of the study system (the non-movement approach; see Bjørneraas et al., 2010). Since it is both difficult and prohibitively time consuming to exactly reproduce expert judgement when dealing with large volumes of tracking data from multiple individuals, some automation is necessary. Users should first manually examine a representative subset of tracks and attempt to visually identify problems — either with individual positions, or with subsets of the track — that persist after filtering on system-generated attributes. Once such problems are identified, users can conceptualise algorithms that can be applied to their data to resolve them.

A common example of a problem with individual positions is that of point outliers or ‘spikes’ (Bjørneraas et al., 2010), where a single position is displaced far from the track (see Fig. 3.b). Point outliers are characterised by artificially high speeds between the outlier and the positions before and after (called incoming and outgoing speed, respectively; Bjørneraas et al., 2010), lending a ‘spiky’ appearance to the track. Removing spikes is simple: remove positions with extreme incoming and outgoing speeds. Users must first define plausible upper limits of the study species’ speed (Calenge et al., 2009; Seidel et al., 2018). Here, it is important to remember that speed estimates are scale-dependent; high-throughput tracking typically overestimates the speed between positions where the animal is stationary or moving slowly, due to small-scale location errors (Noonan et al., 2019; Ranacher et al., 2016). Even after data with large location errors have been removed, it is advisable to begin with a liberal (high) speed threshold that excludes only the most unlikely speeds. Estimates of maximum speed may not always be readily obtained for all species, and an alternative is to use a data-driven threshold such as the 90^th^ percentile of speeds from the track. Once a speed threshold *S* has been chosen, positions with incoming *and* outgoing speeds > *S* may be identified as spikes and removed.

Some species can realistically achieve speeds > *S* in fast transit segments when assisted by their environment, such as birds with tailwinds, and a simple filter on incoming and outgoing speeds would exclude this valid data. To avoid removing valid, fast transit segments while still excluding spikes, the speed filter can be combined with a filter on the turning angles of each position (see Bjørneraas et al., 2010; Calenge et al., 2009). This combined filter assumes that positions in high-throughput tracking with both high speeds and large turning angles are likely to be due to location errors, since most species are unable to turn sharply at very high speed. Users can then remove those positions whose incoming and outgoing speeds are both > `S`, and where *θ* > `A` (sharp, high-speed turns), where *θ* is the turning angle, and `A` is the turning angle threshold. Many other track metrics may be used to identify implausible movement, and used to filter data (Seidel et al., 2018). We implement spike removal using the `atl_filter_covariates` function (Listing 3).

**Listing 3.**
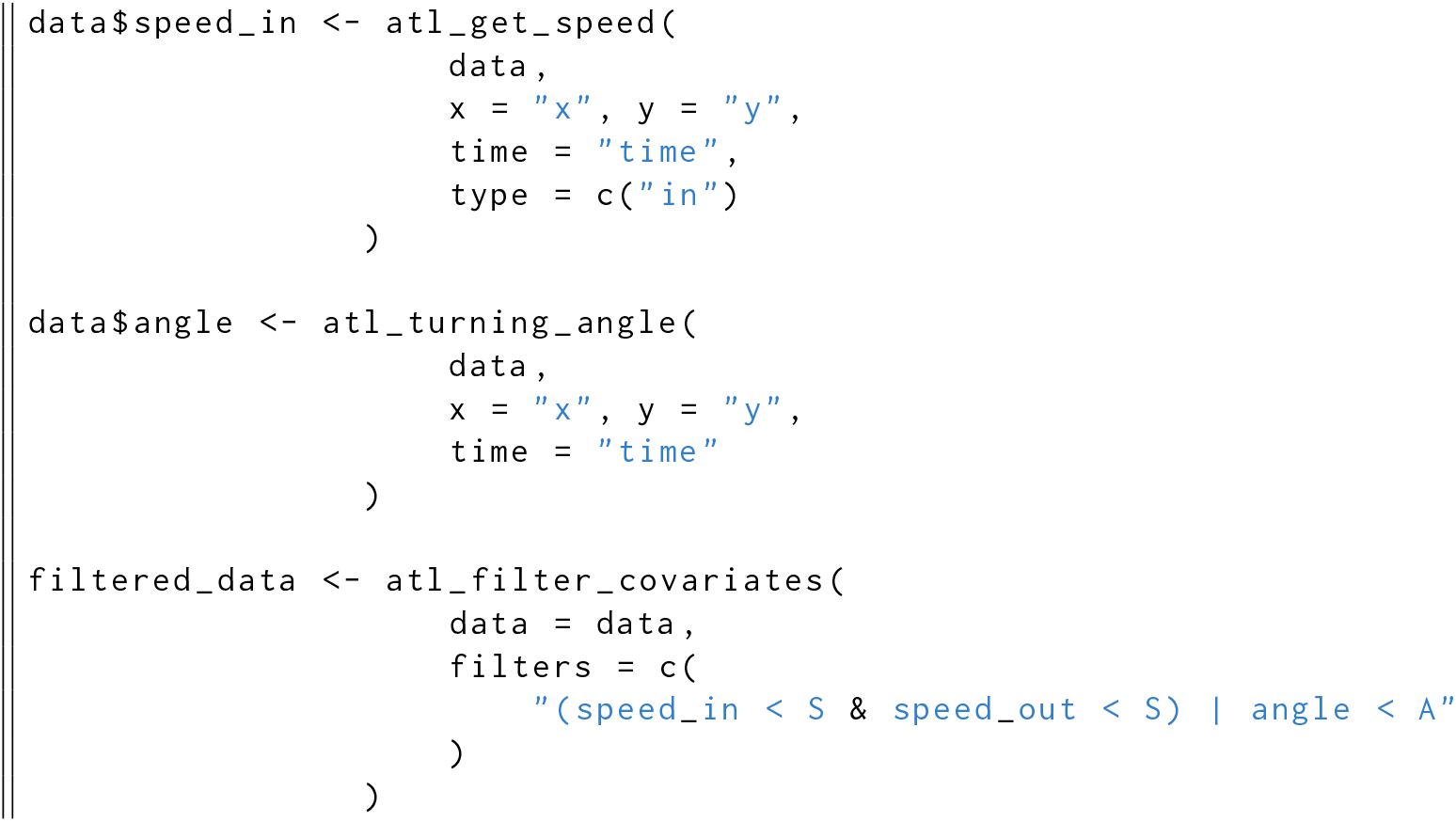
Filtering a movement track on incoming and outgoing speeds, and on turning angle to remove unrealistic movement. The functions `atl_get_speed` and `atl_turning_angle` are used to get the speeds and turning angles before filtering, and assigned to a column in the data (assignment of `speed_out` is not shown). The filter step only retains positions with speeds below the speed threshold *S or* angles above the turning angle threshold *θ*, i.e., positions where the animal is slow but makes sharp turns, and data where the animal moves quickly in a relatively straight line.

Sometimes, entire subsets of the track may be affected by the same large-scale location error. For instance, multiple consecutive positions may be roughly translated (geometrically) away from the real track and form ‘prolonged spikes’, or ‘reflections’ (see Fig. 3.b). These cannot be corrected by targeted removal of individual positions, as in Bjørneraas et al.’s approach (2010), since there are no positions with both high incoming and outgoing speeds, as well as sharp turning angles, that characterise spikes. Since filtering individual positions will not suffice, algorithms to correct such errors must take a track-level view, and target the displaced sequence overall. Track-subset algorithms are likely to be system-specific, and may be challenging to conceptualise or implement. In the case of prolonged spikes, one relatively simple solution is identifying the bounds of displaced segments, and removing positions between them. Users are strongly encouraged to visualise their data before and after applying such algorithms. We caution that these methods are not foolproof, and data that are heavily distorted by errors affecting entire track-subsets should be used with care when making further inferences.

## 6 Smoothing and Thinning Data

### 6.1 Median Smoothing

After filtering out large location errors, the track may still look ‘spiky’ at small scales, and this is due to smaller location errors that are especially noticeable when the individual is stationary or moving slowly (Noonan et al., 2019). These smaller errors are challenging to remove since their attributes (such as speed and turning angles) are within the expected range of movement behaviour for the study species. The large data volumes of high-throughput tracking allow users to resolve this problem by smoothing the positions. The most basic ‘smooths’ work by approximating the value of an observation based on neighbouring values. For a one-dimensional series of observations, the neighbouring values are the *K* observations centred on each index value *i*. The range *i −* (*K −* 1)/2 … *i* + (*K −* 1)/2 is referred to as the moving window as it shifts with *i*, and *K* is the moving window size. A common smooth is nearest neighbour averaging, in which the value of an observation *x_i_* is the average of the moving window *K*. The median smooth is a variant of nearest neighbour averaging which uses the median rather than the mean, and is more robust to outliers (Tukey 1977). The median smoothed value of the X coordinate, for instance, is

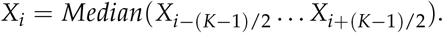

Users can apply a median smooth with an appropriate *K* independently to the X and Y coordinates of a movement track to smooth it (see Fig. 4.a – e). The median smooth is robust to even very large temporal and spatial gaps, and does not interpolate between positions when data are missing. Thus it is not necessary to split the data into segments separated by periods of missing observations when applying the filter (see Fig. 4).

**Figure 4.**
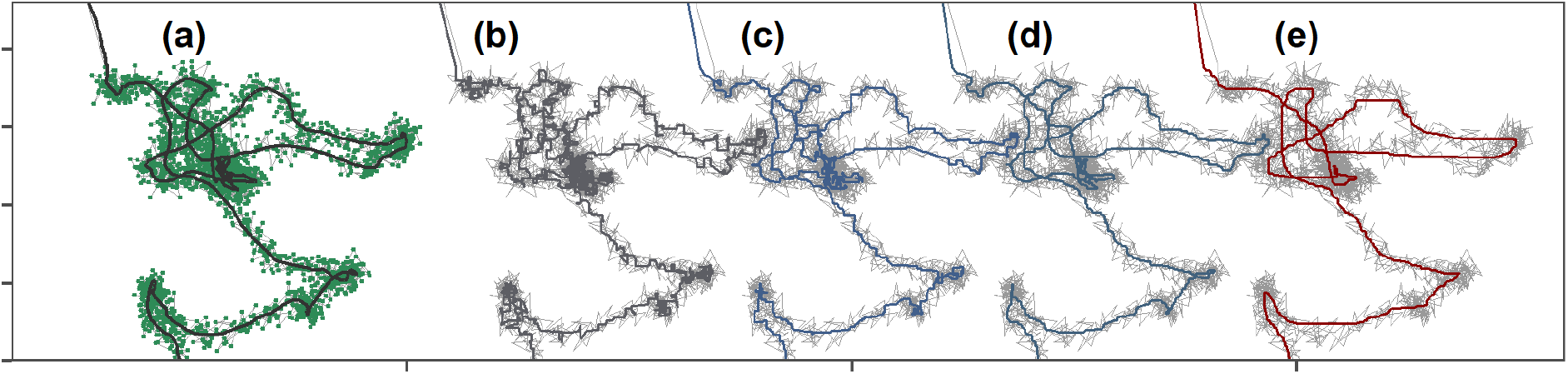
Median smoothing position coordinates reduces small-scale location error in tracking data. The goal of this step is to approximate the simulated canonical track (black line, **(a)**), given positions with small-scale error remaining from previous steps (green points; grey lines). Median smoothing the given position coordinates (green points, in **(a)**) over a moving window (*K*) of **(b)** 5, **(c)** 11, and **(d)** 21 positions respectively, yields approximations (coloured lines) of the canonical track. **(e)** However, extremely large *K* (101) may lead to a loss of both large and small scale detail. Grey lines show the track without smoothing. Users are cautioned that there is no correct *K*, and they must subjectively choose a *K* which most usefully trades small-scale details of the track for large-scale accuracy.

Some data sources, such as GPS, provide tracks that have already been smoothed in quite sophisticated ways, such as with a Kalman filter, making a median smooth unnecessary (Kaplan and Hegarty, 2005). Furthermore, smoothing is not a panacea for data quality issues, and has its drawbacks. Smoothing does not change the number of observations, but does decouple the coordinates from some of their attributes. For instance, smoothing breaks the relationship between a coordinate and the location error estimate around it (VARX, VARY, and SD in ATLAS systems). Since the X and Y coordinates are smoothed independently, the smoothed coordinates of an observation will likely differ from all the coordinates used to compute the smoothed value. Consequently, the location error estimate around each coordinate, and around the localisation overall, become invalid and should be ignored. This makes subsequent filtering on measures of data quality unreliable, and smoothed data are unsuitable for use with methods that model location uncertainty (Calabrese et al., 2016; Fleming et al., 2014, 2020; Noonan et al., 2019). Furthermore, any position covariates (e.g. environmental values such as landcover or elevation) obtained before smoothing should be replaced with the covariates obtained at the smoothed coordinates. Thus, when applying location error modelling methods, users should ensure that the error measure bears a mechanistic relationship with the location estimate (see Fleming et al., 2020; Noonan et al., 2019, for more details). Additionally, excessively large *K* may result in a loss in detail of the individual’s small-scale movement (compare Fig. 4.e with 4.a). Users must themselves judge how best to balance large-scale and small-scale accuracy, and choose *K* accordingly. Median smoothing is provided by the `atlastools` function `atl_median_smooth`, with the only option being the moving window size, which must be an odd integer (Listing 4).

**Listing 4.**
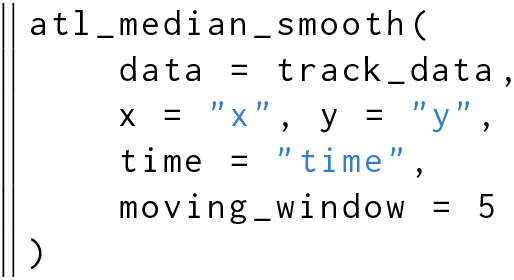
Median smoothing a movement track using the function `atl_median_smooth` function with a moving window *K = 5*. Larger values of *K* yield smoother tracks, but *K* should always be some orders of magnitude lower than the number of observations.

### 6.2 Thinning Movement Tracks

Most data at this stage are technically ‘clean’, yet the volume alone may pose challenges for lower-specification or older hardware and software if these are not optimised for efficient computation. Thinning data i.e., reducing their volume, need not compromise researchers’ ability to answer ecological questions; for instance, proximity-based social interactions lasting 1 – 2 minutes would still be detected on thinning from a sampling interval of 1 second to 1 minute (Aspillaga et al., 2021a). Thinning data also does not imply that efforts to collect high-throughput movement data are ‘wasted’, as rich movement datasets enable more detailed and more accurate representation of the true track, as elaborated above. Indeed, some analyses require that temporal auto-correlation in the data be broken by subsampling the data to a lower resolution; these include traditional kernel density estimators for animal home-range, as well as resource selection functions (Dupke et al., 2017; Fleming et al., 2014; Manly et al., 2007). Furthermore, a number of powerful methods in movement ecology, including Hidden Markov Models and integrated Step-Selection Analysis recommend uniform sampling intervals (Avgar et al., 2016; Langrock et al., 2012; Michelot et al., 2016). Finally, subsampling data may be an important strategy in exploratory data analysis; for instance, it allows researchers to determine whether computationally intensive methods, such as distance and speed estimates from continuous time movement model fitting, are required for their data, or whether the movement metrics stabilise at a certain time scale (Noonan et al., 2019). Two plausible approaches here are subsampling and aggregation, and both approaches begin with identifying time-interval groups (e.g. of 1 minute). Subsampling picks one position from each time-interval group while aggregation involves computing the mean or median of all system-generated attributes for positions within a time-interval group. Both approaches yield one position per time-interval group (Fig. 5.a). Categorical variables, such as the habitat type associated with each position, can be aggregated using a suitable measure such as the mode. We caution users that thinning causes an extensive loss of small-scale detail in the data, and should be used carefully.

**Figure 5.**
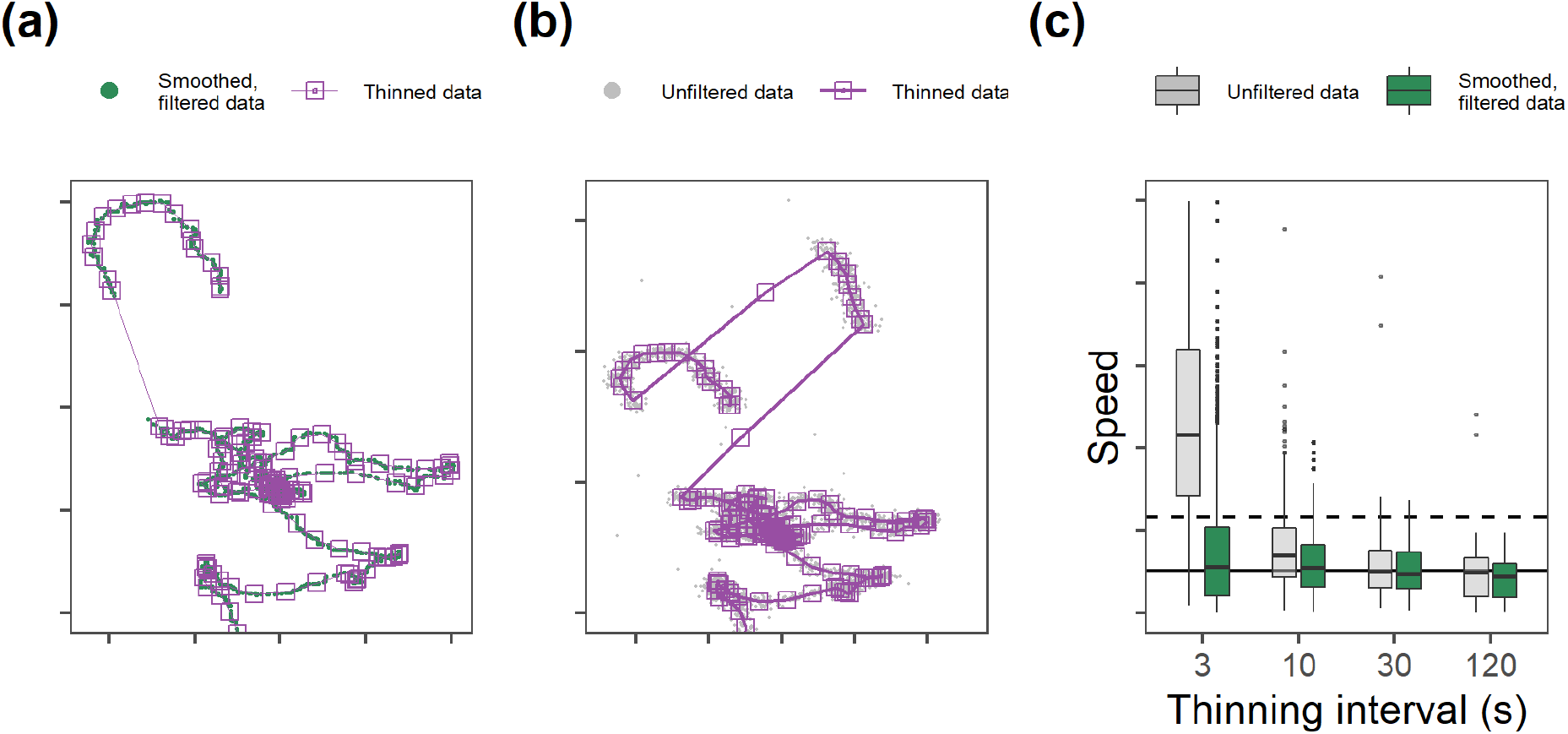
Thinning tracking data can aid computation but must be approached carefully. Aggregating a filtered and smoothed movement track **(a)** preserves track structure while reducing data-volume, but **(b)** aggregating before filtering gross location errors and unrealistic movement leads to the persistence of large-scale errors (such as prolonged spikes). **(c)** Thinning before data cleaning can lead to significant misestimations of essential movement metrics such as speed at lower intervals. Boxplots show the median and interquartile ranges for speed estimates of tracks aggregated over intervals of 3, 10, 30, and 120 seconds. For comparison, the median and 95^th^ percentile of speed of the canonical track are shown as solid and dashed horizontal lines, respectively.

Both aggregation and subsampling have their relative advantages. The aggregation method is less sensitive to selecting point outliers by chance than subsampling. However, to account for location error with methods such as state-space models (Johnson et al., 2008; Jonsen et al., 2005, 2003) or continuous time movement models (Calabrese et al., 2016; Fleming et al., 2014, 2020; Gurarie et al., 2017; Noonan et al., 2019), correctly propagating the location error is important, and subsampling directly propagates these errors without further processing. In the aggregation method, the location error around each coordinate provided by either GPS or reverse-GPS systems can be propagated to the averaged position as the sum of errors divided by the square of the number of observations contributing to each average (*N*):

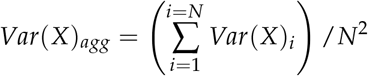

Similarly, the overall location error estimate for the average of *N* positions in a time-interval can be calculated by treating it as a variance. For instance, the ATLAS error and GPS error measures (SD and HDOP, respectively) can be aggregated as:

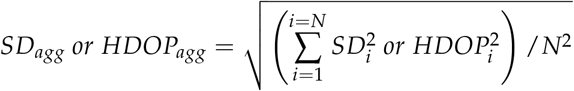

Users may question why thinning, which can obtain consensus positions over an interval and also reduce data-volumes should not be used directly on the raw data. We caution that thinning prior to excluding unrealistic movement and smoothing (Fig 5.b) can lead to preserving artefacts in the data, and estimates of essential metrics — such as straight-line displacement (and hence, speed) — that are substantially different from the true value (see Fig. 5.c; Noonan et al., 2019). In our example, the data with errors would have to be thinned to 1⁄30^th^ of its volume for the median speed of the thinned data to be comparable with the overall median speed — this is an undesirable step if the aim is fine-scale tracking. Additionally, the optimal level of thinning can be difficult to determine, especially if there is wide individual variation in movement behaviour, and the mis-estimation of track metrics from inappropriately thinned data could have consequences for the implementation of subsequent filters based on detecting unrealistic movement. However, thinning before data-cleaning has its place as a useful step before exploratory visualisation of the movement track, since reduced data-volumes are easier to handle for plotting software. Thinning is implemented in `atlastools` using the `atl_thin_data` function, with either aggregation or subsampling (specified by the `method` argument) over an interval using the `interval` argument. Grouping variable names (such as animal identity) may be passed as a character vector to the `id_columns` argument (Listing 5).

**Listing 5.**
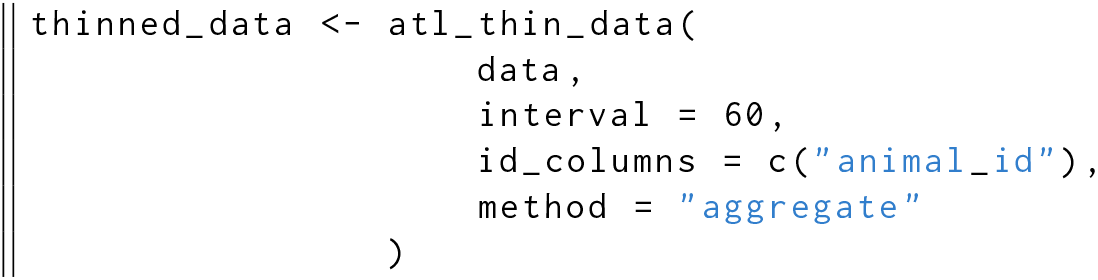
Code to thin data by aggregation in `atlastools`. The method can be either “aggregate” or “subsample”. The time interval is specified in seconds, while the `id_columns` allows a character vector of column names to be passed to the function, with these columns used as identity variables. Both methods return a dataset with one rows per time-interval.

## 7 Creating System-Specific Pre-processing Tools

Researchers’ pre-processing requirements may exceed the functionalities of existing tools, and they will have to conceptualise and implement their own methods. For instance, an important and common analysis with animal tracking data is to link space use with environmental covariates. This is difficult even with smoothed and thinned high-throughput data, as these may be too large for statistical packages, or have strong autocorrelation. Users aiming for such analyses can benefit from segmenting and clustering the data into spatio-temporally independent bouts of different behavioural modes (Patin et al., 2020). Treating these as the unit of observation also conveniently sidesteps pseudo-replication and reduces computational requirements. Common methods of segmentation-clustering are often not scalable to very large or gappy datasets (Langrock et al., 2012; Michelot et al., 2016; Patin et al., 2020), making a first-principles approach an essential part of a pre-processing pipeline, and one which may help in the further implementation of parametric error-modelling approaches (Fleming et al., 2014; Noonan et al., 2019). Here we focus on a simple yet robust segmentation-clustering algorithm to make ‘residence patches’, identified as bouts of relatively stationary behaviour (Barraquand and Benhamou, 2008; Bijleveld et al., 2016; Oudman et al., 2018). Details of the implementation may be found in the package code, and examples are provided in the Supplementary Material.

### 7.1 Conceptualising a Simple Segmentation-Clustering Algorithm: The Residence-Patch Example

Before implementing the algorithm, users should identify positions where the animal is relatively stationary, for instance on its speed or first-passage time (Barraquand and Benhamou, 2008; Bracis et al., 2018). The algorithm should begin by assessing whether consecutive stationary positions are spatio-temporally independent, and clusters them together into a residence patch if they are not. This clustering could be based on a simple proximity threshold — points further apart than some threshold distance are likely to represent two different residence patches. In cases where animals visit multiple sites in sequence (such as traplining Thomson et al., 1997), and which researchers might wish to consider as a single residence patch, a larger-scale distance threshold can help cluster nearby residence patches together, and this can also be applied to cluster together patches artificially separated due to missing data. Similarly, missing data between two consecutive observations at a similar location, but at two very different time points, can be separated by a time-difference threshold, which can also apply to patches that would otherwise be clustered by the large-scale distance threshold. Users are encouraged to base these thresholds on the movement habits of their study species (see the Worked Out Example).

**Listing 6.**
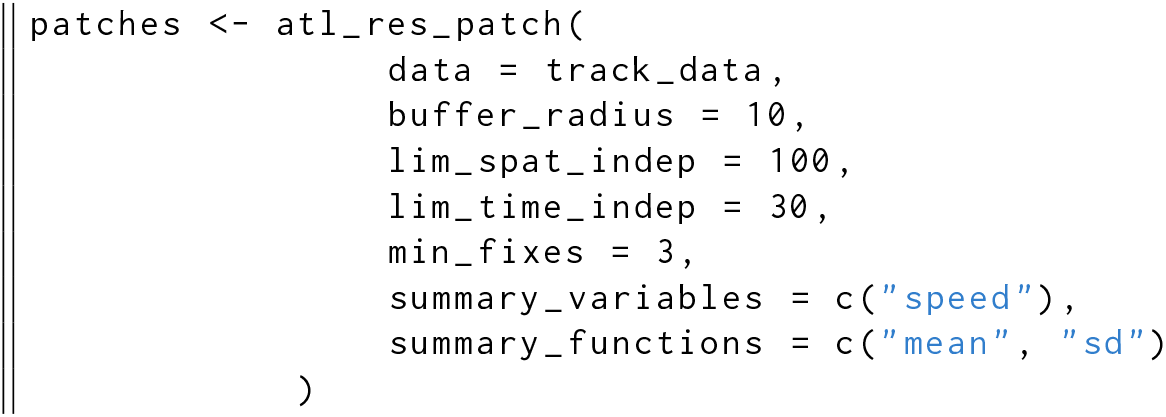
The `atl_res_patch` function can be used to classify a track into residence patches. The arguments buffer_`radius` and `lim_spat_indep` are specified in metres, while the `lim_time_indep` is provided in minutes. In this example, specifying `summary_variables = c(“speed”)`, and `summary_functions = c(“mean”, “sd”)` will provide the mean and standard deviation of instantaneous speed in each residence patch. The `atl_patch_summary` function is used to access the classified patch in one of three ways, here using the summary option which returns a table of patch-wise summary statistics.

We have implemented a working example of the simple clustering concept presented here as the function `atl_res_patch` (see Fig. 6.b; Listing 6), which requires three parameters: (1) the distance threshold between positions (called `buffer_size`), (2) the large-scale distance threshold between clusters of positions (called `lim_spat_indep`), and (3) the time-difference threshold between clusters (called `lim_time_indep`). Clusters formed of fewer than a minimum number of positions can be excluded. Our residence patch algorithm is capable of correctly identifying clusters of related points from a simulated movement track (Fig. 6). This includes clusters where the animal is relatively stationary (orange and green patches, Fig. 6.a), as well as clusters where the animal is moving slowly (blue patch, Fig. 6.a). Such flexibility is especially useful when studying movements that may represent two different modes of the same behaviour, for instance, area-restricted search, as well as slow, searching behaviour with a directional component. It is important to systematically test such custom-made algorithms, to ensure reproducibility (Marwick et al., 2018; Wickham, 2015). Simple examples of such tests for the residence patch and other functions in `atlastools` may be found in the `tests/` directory in the associated Github repository.

**Figure 6.**
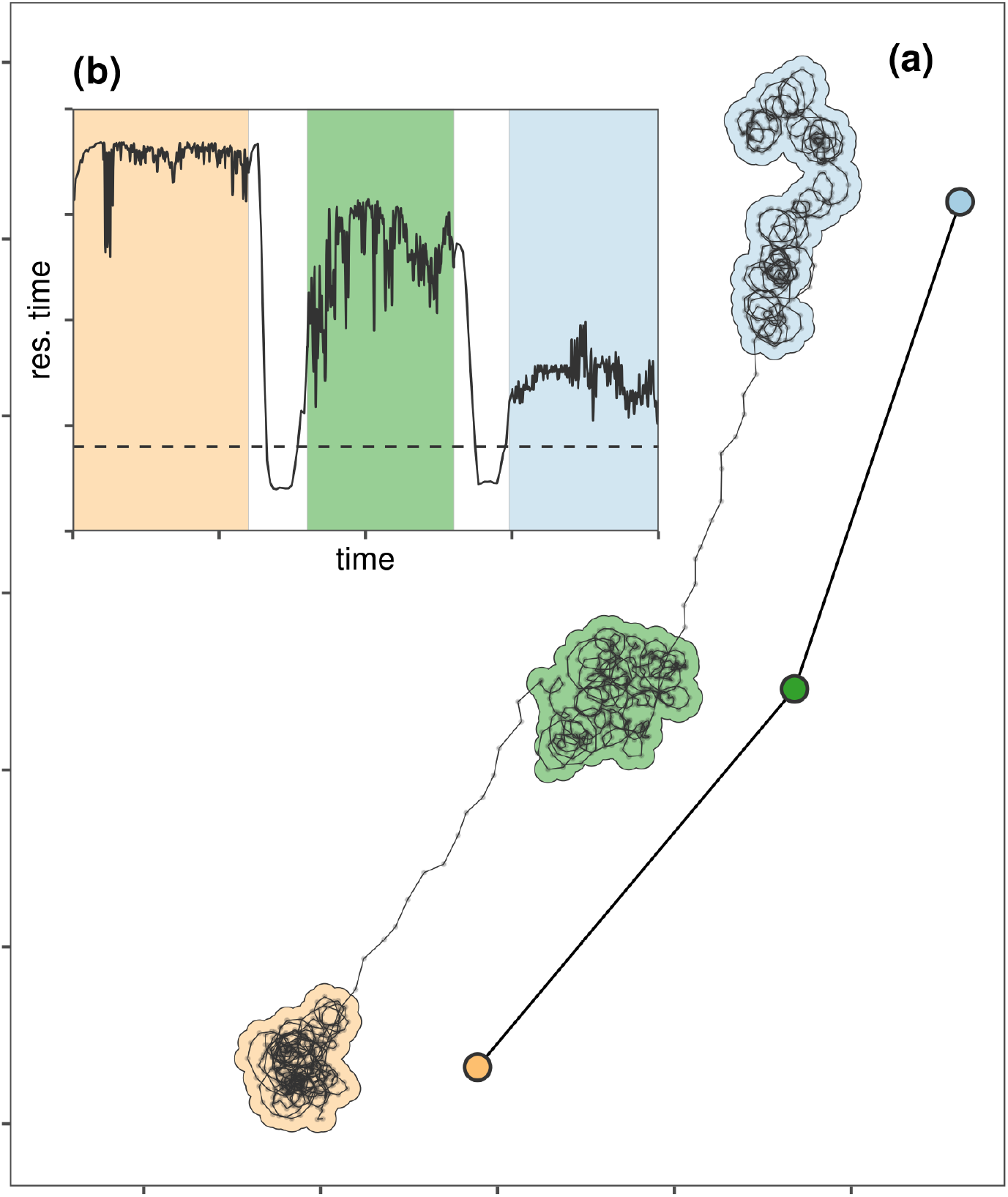
Movement tracks can be classified into residence patches, while leaving out the transit between them. **(a)** The residence patch method correctly identifies clusters of positions where the individual is relatively stationary (orange and green patches), as well as positions where it is moving slowly (blue patch). This is especially useful when studying more complex behaviour such as area-restricted search, which may have a directional component. The residence patch method loses the details of movement between patches, but can efficiently represent the general pattern of space use (see coloured points representing patch centroids, and lines joining them). **(b)** A plot of residence time against time (solid line; Bracis et al. 2018) shows how the residence patch algorithm segments and clusters positions of prolonged residence. Users should remove transit segments beforehand. Regions are shaded by the temporal bounds of each residence patch. The arguments passed to `atl_res_patch` determine the clustering, and can be adjusted to fit the study system. Users are cautioned that there are no ‘correct’ arguments, and the best guide is the biology of the tracked individual.

### 7.2 A Real-World Test of User-Built Pre-Processing Tools

We applied the pre-processing pipeline using `atlastools` functions described above to a calibration dataset to verify that the residence patch method could correctly identify known stopping points (see Fig. 7). We collected the calibration data (n = 50,816) on foot and by boat, with a hand-held WATLAS tag (sampling interval = 1s) around the island of Griend (53.25°N, 5.25°E) in August 2020 (WATLAS: Wadden Sea ATLAS system Beardsworth et al., 2021a, Bijleveld et al. *in prep.*). Stops in the calibration track were recorded as waypoints using a handheld GPS device (Garmin Dakota 10). We estimated the real duration of each stop as the time difference between the first and last position recorded within 50m of each waypoint, within a 10 minute window before and after the waypoint timestamp (to avoid biased durations from revisits). Stops had a median duration of 10.28 minutes (range: 1.75 minutes – 20 minutes; see Supplementary Material). We cleaned the data before constructing residence patches by (1) removing a single outlier (> 15 km away), removing unrealistic movement (≥ 15 m/s), smoothing the data (*K* = 5), and (4) thinning the data by subsampling over a 30 second interval. The cleaning steps retained 37,324 positions (74.45%), while thinning reduced these to 1,803 positions (4.8% positions of the smoothed track). Details and code are provided in the Supplementary Material (see Validating the Residence Patch Method with Calibration Data).

**Figure 7.**
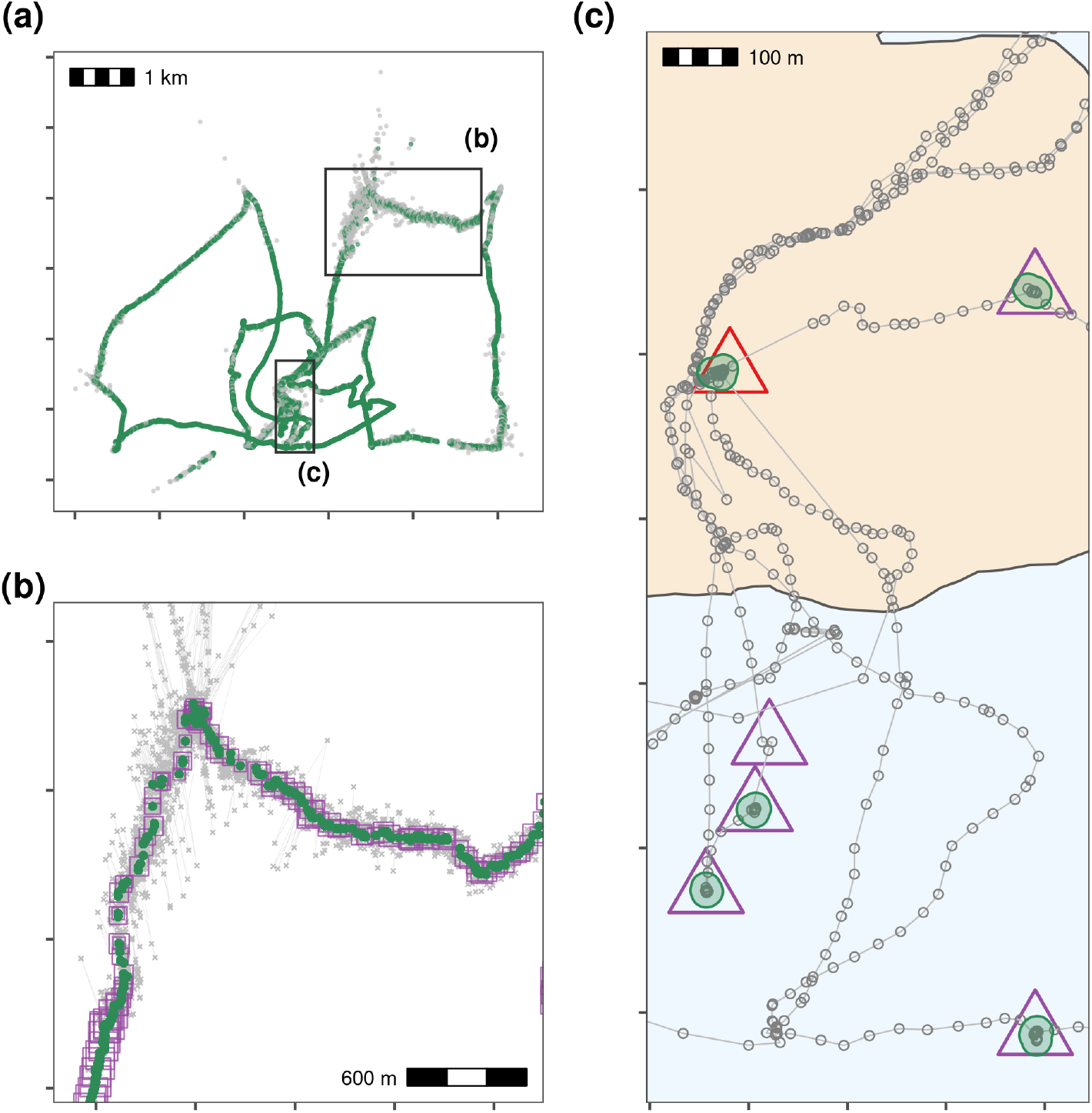
Pre-processing steps for WATLAS calibration data showing filtering on speed, median smoothing and thinning by aggregation, and making residence patches. **(a)** Positions with incoming and outgoing speed > 15 m/s are removed (grey crosses = removed, green points = retained). **(b)** Raw data (grey crosses), median smoothed positions (green circles; moving window *K* = 5), and the smoothed track thinned by aggregation to a 30 second interval (purple squares). Square size corresponds to the number of positions used to calculate the averaged position during thinning. **(c)** Clustering thinned data into residence patches (green polygons) yields robust estimates of the location of known stops (purple triangles). The algorithm identified all areas with prolonged residence, including those which we had not intended to be recorded, such as stops at the field station (n = 12; red triangle). The algorithm also failed to find two stops of 6 and 15 seconds duration, since these were lost in the data thinning step; one of these is shown here (purple triangle without green polygon).

We identified stationary positions as those where the median smoothed speed (*K* = 5) was < 2m/s, as people or a boat moving any faster are likely to be in transit. We clustered these positions into residence patches with a buffer radius of 5m, spatial independence limit of 50m, temporal independence limit of 5 minutes, and a minimum of 3 positions per patch. Inferred residence patches corresponded well to the locations of stops (see Fig. 7.c). However, the residence patch algorithm detected more stops than were logged as waypoints (n = 28, n waypoints = 21). One of these was the field station on Griend where the tag was stored between trips (red triangle, Fig. 7.c). The method also did not detect two stops of 105 and 563 seconds (1.75 and 9.4 minutes) since they were data poor and aggregated away in the thinning step (n positions = 6, 15). To determine whether the residence patch method correctly identified the duration of stops in the calibration track, we first extracted the patch attributes using the function `atl_patch_summary`. We then matched the patches to the waypoints by their median coordinates (rounded to 100 metres). We assigned the inferred duration of the stop as the duration of the spatially matched residence patch. We compared the inferred duration with the real duration using a linear model with the inferred duration as the only predictor of the real duration. Inferred duration was a good predictor of the real duration of a stop (linear model estimate = 1.021, t-value = 12.965, *p* < 0.0001, *R*^2^ = 0.908; see Supplementary Material Fig. 1.7). This translates to a 2% underestimation of the stop duration at a tracking interval of 30 seconds.

## 8 A Worked-Out Example on Animal Tracking Data

We present a fully worked-out example of our pre-processing pipeline and residence patch method using movement data from three Egyptian fruit bats (*Rousettus aegyptiacus*) tracked using the ATLAS system in the Hula Valley, Israel (33.1°N, 35.6°E) (Toledo et al. 2020). Code and data can be found in the Supplementary Material and Zenodo repository (see PROCESSING EGYPTIAN FRUIT BAT TRACKS). Data selected for this example were collected over three nights (5^th^ – 7^th^ May, 2018), with an average of 13,370 positions (SD = 2,173; range = 11,195 – 15,542; interval = 8 seconds) per individual. Plotting the tracks revealed potential location errors (see Fig. 1, see also Supplementary Material Fig. 2.1), reduced by removing observations with ATLAS SD > 20 (see Supplementary Material Section 2.5), and observations calculated using fewer than four base stations, altogether trimming 22% of the raw data (mean positions remaining = 10,447 per individual). Then, we removed unrealistic movement represented by positions with incoming and outgoing speeds > 20 m/s that exceed the maximum flight speed recorded in this species (15 m/s; Tsoar et al. 2011), leaving 10,337 positions per individual on average (98% of previous step). We median smoothed the data with a moving window *K* size = 5, and no observations were lost.

We began the construction of residence patches by finding the residence time within 50 metres of each position; this is the maximal radius of a ‘cloud of points’ around fruit trees (Bracis et al., 2018). Foraging bats repeatedly traverse the same routes (Toledo et al. 2020; Tsoar et al. 2011) and this could artificially inflate the residence time of positions along these routes. To avoid confusing revisits with residence, we limited the summation of residence times at each position to the period until the first departure of 5 minutes or more. Thus, two nearby locations (≤ 50m apart) each visited for one minute at a time, but separated by an interval of some hours would not be clustered together as a residence patch. To focus on bat foraging behaviour, we also excluded data collected before a bat left the daily roost (a cave; Fig. 8a) and after its last return to the roost, by excluding locations between 5AM and 8PM; 23,761 of 31,012 positions remained (76.6%). From these positions, we calculated that between leaving the roost to forage, and returning, bats had a mean residence time at each position of 95.64 minutes (SD = 119.23) — this value is still likely to be biased by some positions at the roost. To determine the true duration of foraging, we opted for a first-principles approach and first selected only locations with a residence time > 5 minutes, reasoning that a flying animal stopping for > 5 minutes at a location should plausibly indicate resource use or another interesting localized behaviour. This step retained 5,736 positions per bat on average (17,208 total), or 72.4% of the nighttime positions. We then constructed residence patches with a buffer distance of 25m, a spatial independence limit of 100m, a temporal independence limit of 30 minutes, and rejected patches with fewer than three positions. These values are meant as examples; users should determine the sensitivity of their results to parameter choices. Excluding residence patches at the roost, bats spent 56.95 minutes at foraging sites (SD = 62.20), and were stationary in particular fruit trees and roosting trees during 83.8% of their foraging time (Fig. 8). Although all three bats roosted at the same cave during the day, they used distinct foraging sites across the study area at night (Fig. 8.a). The lack of overlap in tree use, obtained with the residence patch algorithm, shows that co-roosting bats do not necessarily forage on the same trees. Contrasting the actual movement path with the linear path between residence patches can help reveal details of how animal cognition affects space use (Toledo et al., 2020). Bats tended to show prolonged residence near known food sources (fruit trees), but also where no fruit trees were recorded (Fig. 8.b, 8.c), in line with previous evidence for their use of non-fruiting trees to rest, to handle and digest food, and presumably for social interactions (Tsoar et al., 2010).

**Figure 8.**
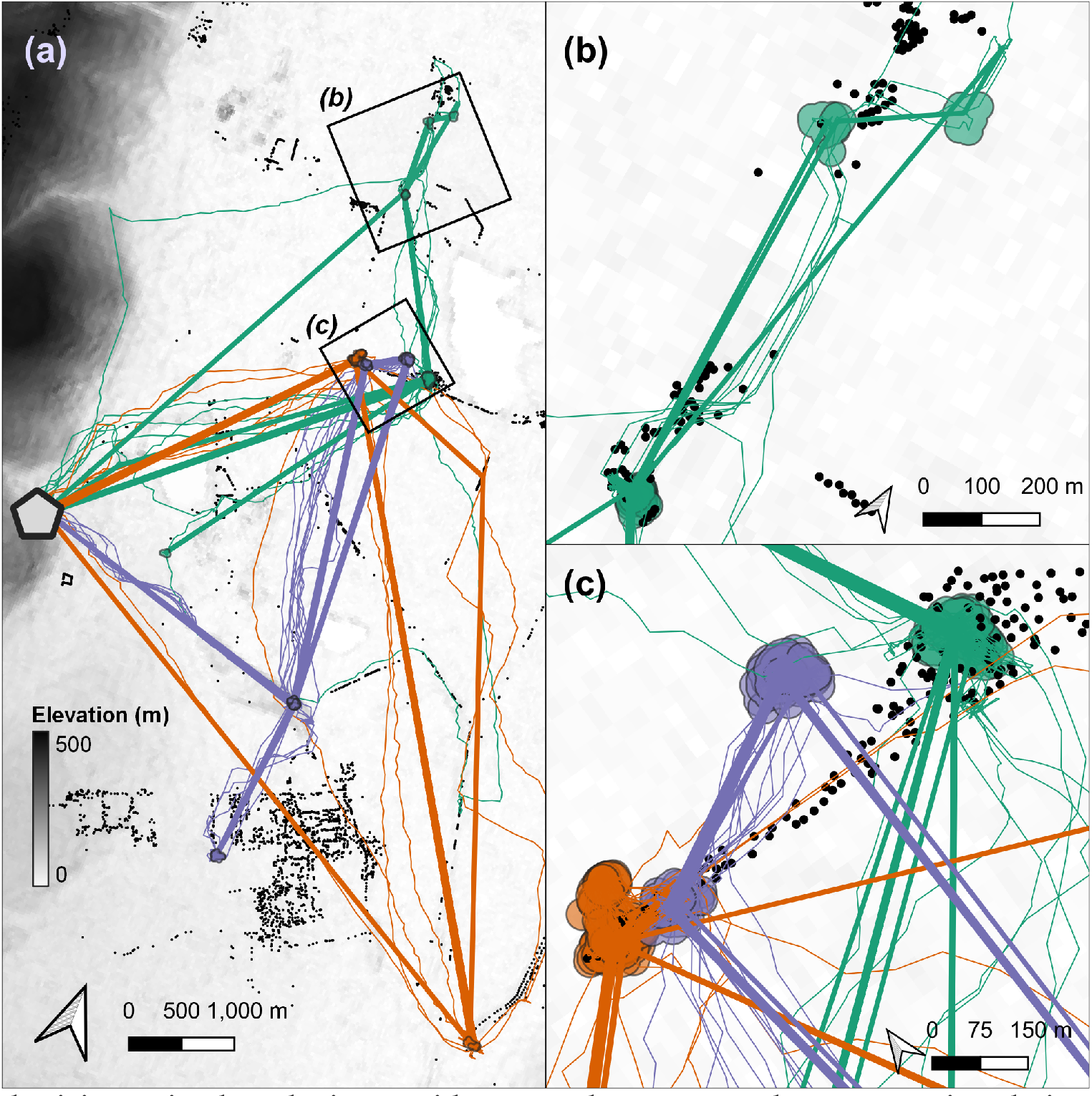
Synthesising animal tracks into residence patches can reveal movement in relation to landscape features, prior exploration, and other individuals. **(a)** Linear approximations of the paths (coloured straight lines) between residence patches (circles) of three Egyptian fruit bats (*Rousettus aegyptiacus*), tracked over three nights in the Hula Valley, Israel. Real bat tracks are are shown as thin lines below the linear approximations, and colours show bat identity. The grey hexagon represents a roost site. Black points represent known fruit trees. Background is shaded by elevation at 30 metre resolution. **(b)** Spatial representations of an individual bat’s residence patches (green polygons) can be used to study site-fidelity by examining overlaps between patches, or to study resource selection by inspecting overlaps with known resources such as fruit trees (black circles). In addition, the linear approximation of movement between patches (straight green lines) can be contrasted with the estimated real path between patches (irregular green lines), for instance, to determine the efficiency of movement between residence patches. **(c)** Fine-scale tracks (thin coloured lines), large-scale movement (thick lines), residence patch polygons, and fruit tree locations show how high-throughput data can be used to study movement across scales. Patches and lines are coloured by bat identity.

## 9 Discussion and Perspective

Recent technical advances in wildlife tracking have already yielded exciting new insights from massive high-resolution movement datasets (Aspillaga et al., 2021a,b; Baktoft et al., 2019, 2017; Beardsworth et al., 2021b,c; Corl et al., 2020; Harel et al., 2016; Harel and Nathan, 2018; Oudman et al., 2018; Papageorgiou et al., 2019; Strandburg-Peshkin et al., 2015; Toledo et al., 2020; Tsoar et al., 2011; Vilk et al., 2021), and high-throughput animal tracking is expected to become increasingly more common in the near future. Tackling the very large datasets that high-throughput tracking generates requires a different approach from that used for traditionally smaller volumes of data. We foresee that movement ecologists will have to adopt ever more practices from fields accustomed to dealing with ‘big data’, and that the field will become increasingly computational (Peng, 2011). Researchers have been informally using some of these approaches, such EDA on small subsets before applying methods to the full data, using efficient tools, and basic batch-processing. Yet formally prescribing these steps can help practitioners avoid pitfalls and implement techniques that make their analyses quicker and more reliable. Standardised principles, implemented a basic pipeline, for approaching data cleaning promote reproducibility across studies, making comparative inferences more robust. The open-source `R` package `atlastools` serves as a starting point for methodological collaboration among movement ecologists, and as a simple working example on which researchers may wish to model their own tools. Efficient location error modelling approaches (Aspillaga et al., 2021b; Fleming et al., 2020) may eventually make data-cleaning optional. Yet cleaning tracking data even partially before modelling location error is faster than error-modelling on the full data, and the removal of large location errors may improve model fits. Thus we see our pipeline as complementary to these approaches (Fleming et al., 2014, 2020). Finally, we recognise that the diversity and complexity of animal movement and data collection techniques often requires system-specific, even bespoke, pre-processing solutions. Though the principles outlined here are readily generalised, users’ requirements will eventually exceed the particular tools we provide. For instance, relational databases are the standard for storing very large datasets, and extending pre-processing pipelines to deal with various data sources is relatively simple, as we show in our Supplementary Material. We see the diversity of animal tracking datasets and studies as an incentive for more users to be involved in developing methods for their systems. We offer our approach to large tracking datasets, and our pipeline and package as a foundation for system-specific tools in the belief that simple, robust concepts are key to methods development that balances system-specificity and broad applicability.

## 10.2 Acknowledgements

PRG would like to thank Pedro M. Santos Neves for introducing PRG to R package development, for help with setting up `atlastools`, and for help with archiving it on Zenodo; Geert Aarts, Jacob RL Gismann, Evy Gobbens, and Roos Kentie for feedback that improved the manuscript; members of the Modelling Adaptive Response Mechanisms Group (Weissing Lab), and the Theoretical Biology department at the University of Groningen for helpful discussions on `atlastools` and the manuscript. We thank the many volunteers, students, and NIOZ staff involved in operating the WATLAS tracking system, and most importantly Frank van Maarseveen, Bas Denissen and Anne Dekinga. We also thank Yotam Orchan, Yoav Bartan, Sivan Margalit, Anat Levi, David Shohami, Ohad Vilk and other members of the Minerva Center for Movement Ecology for their valuable support, and especially the attendees of ATLAS workshops held in May and June 2020 at the Hebrew University of Jerusalem for helpful comments on the pipeline and `atlastools`. Finally, we thank the three reviewer’s whose comments improved this manuscript. This work was partly funded by the Dutch Research Council grant VI.Veni.192.051 awarded to AIB. ATLAS development was funded by the Minerva Foundation grant and the Adelina and Massimo Della Pergola Professor of Life Sciences to RN, and by the Israel Science Foundation grant (ISF ISF-965/15) to RN and ST. PRG was supported by an Adaptive Life Programme grant in the Weissing Lab, made possible by the University of Groningen’s Faculty of Science and Engineering, and the Groningen Institute for Evolutionary Life Sciences (GELIFES).

## 10.3 Authors’ Contributions

PRG wrote the manuscript and inline code snippets, performed the analyses, prepared the figures, and developed the `R` package `atlastools`. CEB and AIB collected the calibration track, and EL collected the bat movement data, roost and fruit tree locations. RN conceived the idea of writing this manuscript, and PRG, AIB, OS, CEB, ST, and EL contributed to its design, and the design of `atlastools`. All authors contributed to the writing of the manuscript, and the design of figures.

## 10.4 Data Availability

The data and source code to reproduce the figures and analyses in this article and in the Supplementary Material can be found in the Zenodo repository at https://doi.org/10.5281/zenodo.4287462.

## 10.5 Supplementary Material

Code and details for worked out examples on calibration data from the Dutch Wadden Sea, and bat tracking data from the Hula Valley, Israel, is provided in the file Supplementary Material 01. This code can also be found (and adapted from) the Github repository: github.com/pratikunterwegs/atlas-best-practices. The reference manual for the `R` package `atlastools` is provided as Supplementary Material 02, and the source code can be found at github.com/pratikunterwegs/atlastools. These materials are also archived in the Zenodo data repository.

