## Supplementary Material 1 for "A Guide to Pre-Processing High-Throughput Animal Tracking Data"

2021-07-01

**3 References** **35**

### 1 Validating the Residence Patch Method with Calibration Data

Here we show how the residence patch method (Barraquand and Benhamou 2008; Bijleveld et al. 2016; Oudman et al. 2018) accurately estimates the duration of known stops in a track collected as part of a calibration exercise in the Wadden Sea. These data can be accessed from the `data` folder at this link: https://doi.org/10.5281/zenodo.4287462. These data are more fully reported in (Beardsworth et al. 2021).

#### 1.1 Outline of Cleaning Steps

We begin by preparing the libraries we need, and installing `atlastools` from Github. After installing `atlastools`, we visualise the data to check for location errors, and find a single outlier position approx. 15km away from the study area (Fig. 1.1, 1.2). This outlier is removed by filtering data by the X coordinate bounds using the function `atl_filter_bounds`; X coordinate bounds $\leq$ 645,000 in the UTM 31N coordinate reference system were removed (n = 1; remaining positions = 50,815; Fig. 1.2). We then calculate the incoming and outgoing speed, as well as the turning angle at each position using the functions `atl_get_speed` and `atl_turning_angle` respectively, as a precursor to targeting large-scale location errors in the form of point outliers. We use the function `atl_filter_covariates` to remove positions with incoming and outgoing speeds $\geq$ the speed threshold of 15 m/s (n = 13,491, 26.5%; remaining positions = 37,324, 73.5%; Fig. 1.3; main text Fig. 7.b). This speed threshold is chosen as the fastest boat speed during the experiment, 15 m/s. Finally, we target small-scale location errors by applying a median smoother with a moving window size $K = 5$ using the function `atl_median_smooth` (Fig. 1.4; main text Fig. 7.c). Smoothing does not reduce the number of positions. We thin the data to a 30 second interval leaving 1,803 positions (4.8% positions of the smoothed track)

#### 1.2 Install `atlastools` from Github

`atlastools` is available from Github and is archived on Zenodo (Gupte 2020). It can be installed using `remotes` or `devtools`. Here we use the `remotes` function `install_github`.

```
if (!require(remotes)) {
  install.packages("remotes", repos = "http://cran.us.r-project.org")
}

# installation using remotes
remotes::install_github("pratikunterwegs/atlastools")
```

#### A Note on :=

The atlastools package is based on data.table, to be fast and efficient (Dowle and Srinivasan 2020). A key feature is modification in place, where data is changed without making a copy. This is already implemented in R and will be familiar to many users as data_frame$column_name <- values.

The data.table way of writing this assignment would be data_frame[, column_name := values]. We use this syntax throughout, as it provides many useful shortcuts, such as multiple assignment:

```
data_frame[, c("col_a", "col_b") := list(values_a, values_b)]
```

Users can use this special syntax, and will find it convenient with practice, but there are *no* cases where users *must* use the data.table syntax, and can simply treat the data as a regular data.frame. However, users are advised to convert their data.frame to a data.table using the function data.table::setDT().

#### 1.3 Prepare libraries

First we prepare the libraries we need. Libraries can be installed from CRAN if necessary.

```
# for data handling
library(data.table)
library(atlastools)
library(stringi)

# for recursion analysis
library(recurse)

# for plotting
library(ggplot2)
library(patchwork)

# making a colour palette
pal <- RColorBrewer::brewer.pal(5, "Set1")
pal[3] <- "seagreen"
```

#### 1.4 Access data and preliminary visualisation

First we access the data from a local file using the data.table package (Dowle and Srinivasan 2020).

In all, we aim to keep three versions of the data: (1) data_raw, the entirely unprocessed data, (2) data, the working version, and (3) data_unproc, data that has been partially processed, but which is one step behind data. This allows us to better illustrate the pre-processing steps, and prevents us from irreversibly modifying our data — at best, we would have to re-run many pre-processing steps, and at worse, we might overwrite the data on disk.

We look at the first few rows, using head(). We then visualise the raw data.

```r
# read and plot example data
data <- fread("data/atlas1060_allTrials_annotated.csv")
data_raw <- copy(data)

# see raw data
head(data_raw)
#>             TAG          TIME NBS VARX VARY COVXY   SD           Timestamp
#> 1: 31001001060 1598027365845   6 6.28 2.85 1.682 3.53 2020-08-21 17:29:25
#> 2: 31001001060 1598027366845   6 2.23 2.23 0.277 2.24 2020-08-21 17:29:26
#> 3: 31001001060 1598027367845   6 2.94 2.82 0.612 2.64 2020-08-21 17:29:27
#> 4: 31001001060 1598027368845   6 8.45 3.68 2.734 4.20 2020-08-21 17:29:28
#> 5: 31001001060 1598027369845   5 6.80 3.26 2.273 3.82 2020-08-21 17:29:29
#> 6: 31001001060 1598027370845   6 3.95 2.94 0.983 2.98 2020-08-21 17:29:30
#>             id      x       y Long  Lat             UTCtime    tID
#> 1: 2020-08-21 650083 5902624 5.25 53.3 2020-08-21 16:29:25 DELETE
#> 2: 2020-08-21 650083 5902624 5.25 53.3 2020-08-21 16:29:26 DELETE
#> 3: 2020-08-21 650073 5902622 5.25 53.3 2020-08-21 16:29:27 DELETE
#> 4: 2020-08-21 650079 5902625 5.25 53.3 2020-08-21 16:29:28 DELETE
#> 5: 2020-08-21 650067 5902621 5.25 53.3 2020-08-21 16:29:29 DELETE
#> 6: 2020-08-21 650071 5902621 5.25 53.3 2020-08-21 16:29:30 DELETE
```

Here we show how data can be easily visualised using the popular plotting package ggplot2. Note that we plot both the points (`geom_point`) and the inferred path between them (`geom_path`), and specify a geospatial coordinate system in metres, suitable for the Dutch Wadden Sea (UTM 31N; ESPG code:32631; `coord_sf`). We save the output to file for future reference.

Since plot code can become very lengthy and complicated, we omit showing further plot code in versions of this document rendered as PDF or HTML; it can however be seen in the online `.Rmd` version.

```r
# plot data
fig_data_raw <-
  ggplot(data) +
  geom_path(aes(x, y),
    col = "grey", alpha = 1, size = 0.2
  ) +
  geom_point(aes(x, y),
    col = "grey", alpha = 0.2, size = 0.2
  ) +
  ggthemes::theme_few() +
  theme(
    axis.title = element_blank(),
    axis.text = element_blank()
  ) +
  coord_sf(crs = 32631)

# save figure
ggsave(fig_data_raw,
  filename = "supplement/figures/fig_calibration_raw.png",
  width = 185 / 25
)
```

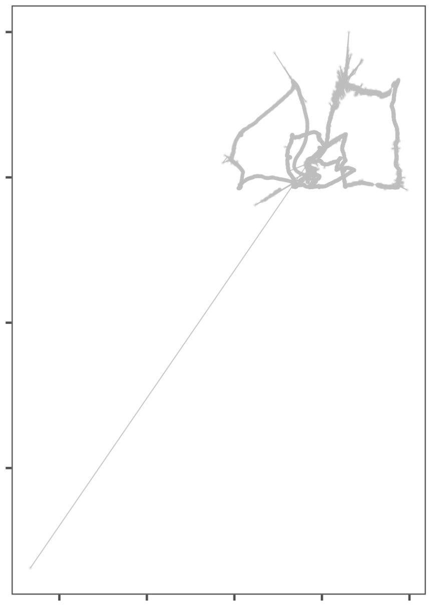

Figure 1.1: The raw data from a calibration exercise conducted around the island of Griend in the Dutch Wadden Sea. A handheld WATLAS tag was used to examine how ATLAS data compared to GPS tracks, and we use the WATLAS data here to demonstrate the basics of the pre-processing pipeline, as well as validate the residence patch method. It is immediately clear from the figure that the track shows location errors, both in the form of point outliers as well as small-scale errors around the true location.

#### 1.5 Filter by bounding box

We first save a copy of the data, so that we can plot the unprocessed data with the cleaned data plotted over it for comparison. Here, `data_unproc`, `data`, and `data_raw` are still the same, since no pre-processing steps have been applied yet.

```
# make a copy using the data.table copy function
data_unproc <- copy(data)
```

We then filter by a bounding box in order to remove the point outlier to the far south east of the main track. We use the `atl_filter_bounds` functions using the `x_range` argument, to which we pass the limit in the UTM 31N coordinate reference system. This limit is used to exclude all points with an X coordinate < 645,000.

We then plot the result of filtering, with the excluded point in black, and the points that are retained in green. After this stage, `data` is filtered and 'ahead' of `data_raw` and `data_unproc`, which are still the same. This pattern will repeat throughout this material.

```
# remove inside must be set to falses
data <- atl_filter_bounds(
  data = data,
  x = "x", y = "y",
  x_range = c(645000, max(data$x)),
  remove_inside = FALSE
)
```

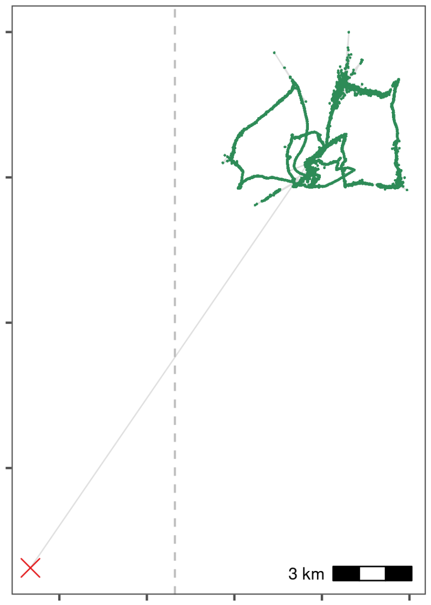

Figure 1.2: Removal of a point outlier using the function `atl_filter_bounds`. The point outlier (black point) is removed based on its X coordinate value, with the data filtered to exclude positions with an X coordinate < 645,000 in the UTM 31N coordinate system. Positions that are retained are shown in green.

#### 1.6 Filter trajectories

##### 1.6.1 Handle time

Time in ATLAS tracks is represented by 64-bit integers (type `long`) that specify time in milliseconds, starting from the beginning of 1970 (the UNIX epoch). This representation of time is called `POSIX` time and is usually specified in seconds, not milliseconds.

Since about 1.6 billion seconds have passed since the beginning of 1970, current `POSIX` times in milliseconds cannot be represented by R's built-in 32-bit integers. A naive conversion results in truncation of out-of-range numbers leading to huge errors (dates many thousands of years in the future).

R does not natively support 64-bit integers. One option is to use the bit64 package, which adds 64-bit integer support to R.

A simpler solution is to convert the times to R's built in `double` data type (also called `numeric`), which uses a 64-bit floating point representation. This representation can represent integers with up to 16 digits without error; we only need 13 digits to represent the number of milliseconds since 1970, so the conversion is error free. We can also perform the conversion and then divide by 1000 so that times are represented in seconds, not milliseconds; this simplifies speed estimation.

If second-resolution is accurate enough (it is for our purposes), the solution that we use is to divide times by 1000 to reduce the resolution from milliseconds to seconds and then to convert the time stamps to R integers. In the spirit of not destroying data, we create a second lower-case column called `time` to store this

```
# divide by 1000, convert to integer, then convert to POSIXct
```

```
data[, time := as.integer(
  as.numeric(TIME) / 1000
)]
```

##### 1.6.2 Add speed and turning angle

```
# add incoming and outgoing speed
data[, `:=`(
  speed_in = atl_get_speed(data,
    x = "x",
    y = "y",
    time = "time"
  ),
  speed_out = atl_get_speed(data, type = "out")
)]

# add turning angle
data[, angle := atl_turning_angle(data = data)]
```

##### 1.6.3 Get 90th percentile of speed and angle

```
# use sapply
speed_angle_thresholds <-
  sapply(data[, list(speed_in, speed_out, angle)],
    quantile,
    probs = 0.9, na.rm = T
  )
```

##### 1.6.4 Filter on speed

Here we use a speed threshold of 15 m/s, the fastest known boat speed. We then plot the data with the extreme speeds shown in grey, and the positions retained shown in green.

Here, `data_unproc` moves 'ahead' of `data_raw`, and holds the data filtered by a bounding box — `data` is also moving ahead, and will be filtered on speed.

```
# make a copy
data_unproc <- copy(data)

# remove speed outliers
data <- atl_filter_covariates(
  data = data,
  filters = c("(speed_in < 15 & speed_out < 15)")
)

# recalculate speed and angle
data[, `:=`(
  speed_in = atl_get_speed(data,
    x = "x",
    y = "y",
```

```
    time = "time"
  ),
  speed_out = atl_get_speed(data, type = "out")
)]

# add turning angle
data[, angle := atl_turning_angle(data = data)]
```

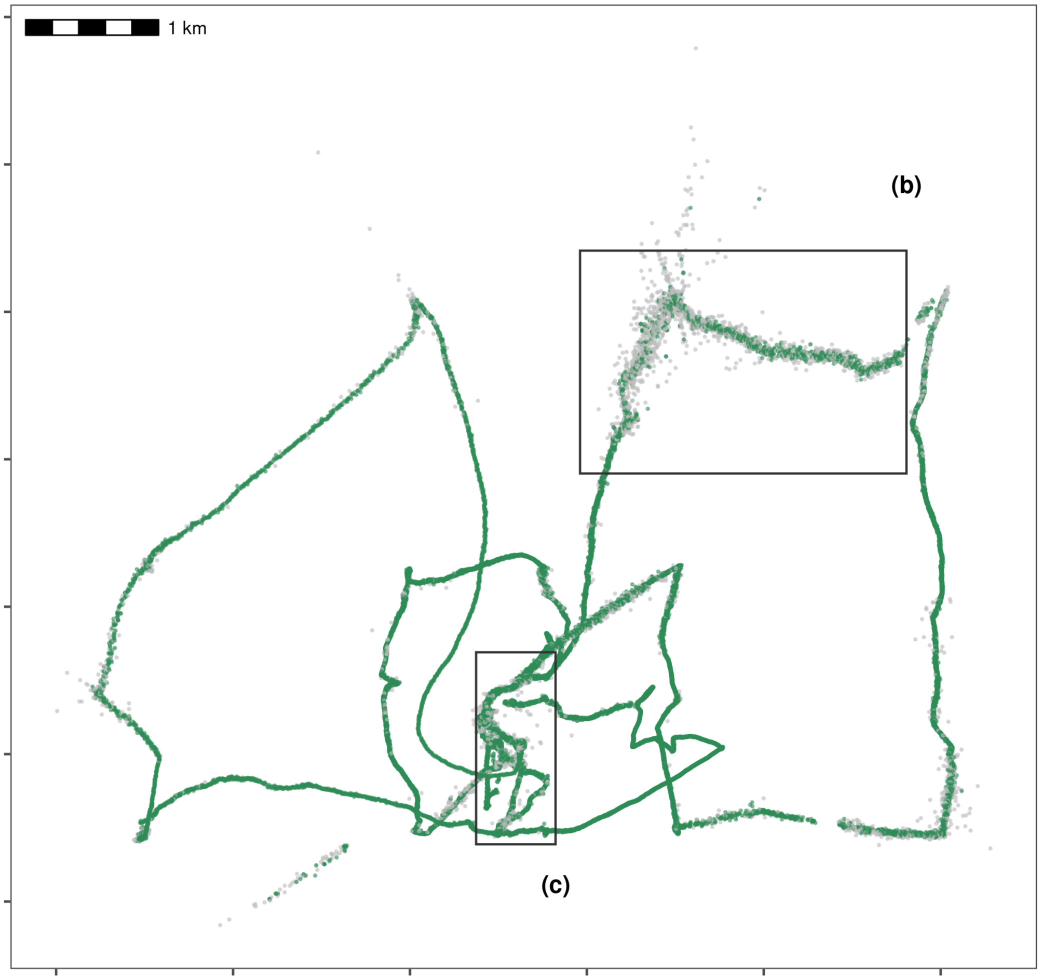

Figure 1.3: Improving data quality by filtering out positions that would require unrealistic movement. We removed positions with speeds $\geq$ 15 m/s, which is the fastest possible speed in this calibration data, part of which was collected in a moving boat around Griend. Grey positions are removed, while green positions are retained. Rectangles indicate areas expanded for visualisation in following figures.

#### 1.7 Smoothing the trajectory

We then apply a median smooth over a moving window ($K$ = 5). This function modifies in place, and does not need to be assigned to a new variable. We create a copy of the data before applying the smooth so that we can compare the data before and after smoothing.

```
# apply a 5 point median smooth, first make a copy
data_unproc <- copy(data)

# now apply the smooth
atl_median_smooth(
  data = data,
  x = "x", y = "y", time = "time",
  moving_window = 5
)
```

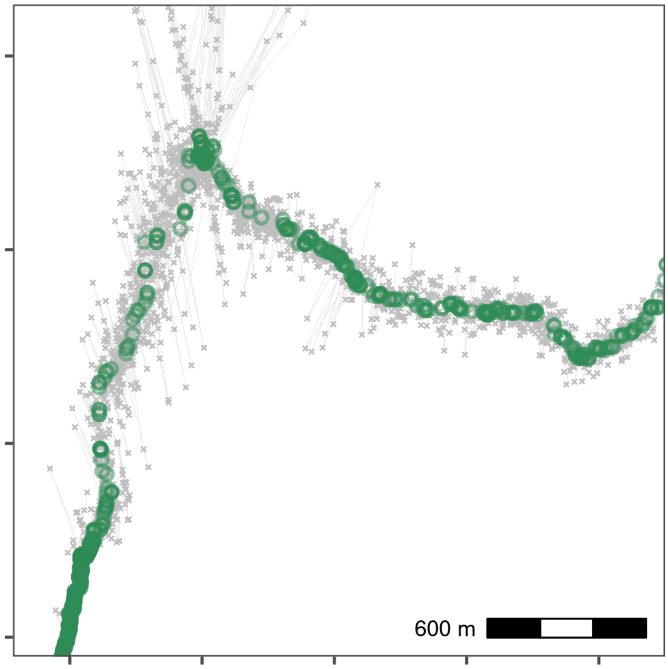

Figure 1.4: Reducing small-scale location error using a median smooth with a moving window $K = 5$. Median smoothed positions are shown in green, while raw, unfiltered data is shown in grey. Median smoothing successfully recovers the likely path of the track without a loss of data. The area shown is the upper rectangle from Fig. 1.3.

#### 1.8 Thinning the data

Next we thin the data by aggregation to demonstrate thinning after median smoothing. Following this, we plot the median smooth and thinning by aggregation.

```
# save a copy
data_unproc <- copy(data)

# remove columns we don't need
data <- data[, !c("tID", "Timestamp", "id", "TIME", "UTCtime")]

# thin to a 30s interval
data_thin <- atl_thin_data(
```

```
  data = data,
  interval = 30,
  method = "aggregate",
  id_columns = "TAG"
)
```

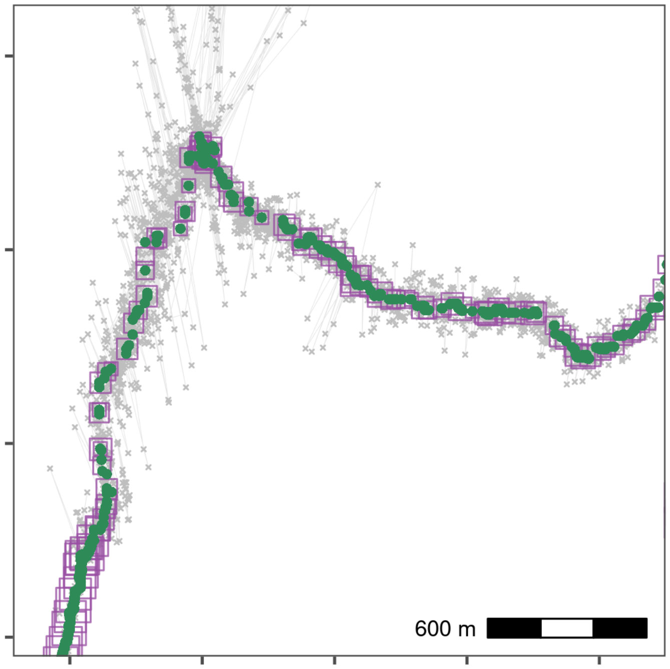

Figure 1.5: Thinning by aggregation over a 30 second interval (down from 1 second) preserves track structure while reducing the data volume for computation. Here, thinned positions are shown as purple squares, with the size of the square indicating the number of positions within the 30 second bin used to obtain the average position. Green points show the median smoothed data from Fig. 1.4, while the raw data are shown in grey. The area shown is the upper rectangle in Fig. 1.3.

#### 1.9 Residence patches

##### 1.9.1 Get waypoint centroids

We subset the annotated calibration data to select the waypoints and the positions around them which are supposed to be the locations of known stops. Since each stop was supposed to be 5 minutes long, there are multiple points in each known stop.

```
data_res <- data_unproc[stri_detect(tID, regex = "(WP)")]
```

From this data, we get the centroid of known stops, and determine the time difference between the first and last point within 50 metres, and within 10 minutes of the waypoint positions' median time.

Essentially, this means that the maximum duration of a stop can be 20 minutes, and stops above this duration are not expected.

```r
# get centroid
data_res_summary <- data_res[, list(
  nfixes_real = .N,
  x_median = median(x),
  y_median = median(y),
  t_median = median(time)
),
by = "tID"
]

# now get times 10 mins before and after
data_res_summary[, c("t_min", "t_max") := list(
  t_median - (10 * 60),
  t_median + (10 * 60)
)]

# manually get the duration of the stops
wp_data <- mapply(function(l, u, mx, my) {

  # first select all data whose timestamp is between
  # the upper and lower bounds of the stop (l = lower, u = upper)
  tmp_data <- data_unproc[inrange(time, l, u), ]

  # calculate the distance between the positions selected above
  # and the median X and Y coordinates of the stop (centroid)
  tmp_data[, distance := sqrt((mx - x)^2 + (my - y)^2)]

  # keep positions that are within 50m of the centroid
  tmp_data <- tmp_data[distance <= 50, ]

  # get the duration of the stop as the difference between
  # the minimum and maximum times of the positions retained above
  return(diff(range(tmp_data$time)))
}, data_res_summary$t_min, data_res_summary$t_max,
data_res_summary$x_median, data_res_summary$y_median,

# this specifies that a vector, rather than a list, is returned
SIMPLIFY = TRUE
)

# get waypoint summary --- rounding median coordinates to the nearest 100m
patch_summary_real <- data_res_summary[, list(
  nfixes_real = nfixes_real,
  x_median = round(median(x_median), digits = -2),
  y_median = round(median(y_median), digits = -2)
),
by = "tID"
]

# add real duration
patch_summary_real[, duration_real := wp_data]
```

##### 1.9.2 Prepare data

First, we filter data where we know the animal (or in this case, the human-carried tag) spent some time at or near a position, as this is the first step to identify residence patches. One way of doing this is by filtering out positions with speeds above which the tag (ideally on an animal) is likely to be in transit. Rather than filtering on instantaneous speed estimates, filtering on a median smoothed speed estimate is more reliable.

##### 1.9.3 Exclude transit points

Here, we aim to remove locations where the tag is clearly moving, by filtering on smoothed speed, using a one-way median smooth with $K = 5$. The speeds between points must be recalculated here because the speed metrics now associated with the data refer to the raw data before median smoothing.

```
# get 4 column data
data_for_patch <- copy(data_thin)

# recalculate speeds, removing speed out
data_for_patch[, c("speed_in", "speed_out") := list(
  atl_get_speed(data_for_patch),
  NULL
)]

# get smoothed speed
data_for_patch[, speed_smooth := runmed(speed_in, k = 5)]

# save recurse data
fwrite(data_for_patch, file = "data/data_calib_for_patch.csv")
```

##### 1.9.4 Run residence patch method

We subset data with a smoothed speed < 2 m/s in order to construct residence patches. From this subset, we construct residence patches using the parameters: `buffer_radius` = 5 metres, `lim_spat_indep` = 50 metres, `lim_time_indep` = 5 minutes, and `min_fixes` = 3.

```
# assign id as tag
data_for_patch[, id := as.character(TAG)]

# on known residence points
patch_res_known <- atl_res_patch(
  data = data_for_patch[speed_smooth < 2, ],
  buffer_radius = 5,
  lim_spat_indep = 50,
  lim_time_indep = 5,
  min_fixes = 3
)
```

##### A note on summary statistics

Users specifying a `summary_variable` should make sure that the variable for which they want a summary statistic is present in the data. For instance, requesting mean speed by passing `summary_variable = "speed"` and `summary_function = "mean"` to `atl_res_patch`, should make sure that their data includes a column called `speed`.

##### 1.9.5 Get spatial and summary objects

Having classified slow-moving or stationary behavioural bouts into residence patches, many animal ecologists would most probably wish to know something about the environment at or around these patches — more accurately, around the point locations classified into patches.

How exactly this is done depends on the relative spatial scales of the residence patches and the resolution of the environmental data layer. For instance, a residence patch some 40m – 50m wide or long may be overlaid on an environmental raster layer with a resolution of 250m. In this case, sampling the layer at the centroid of the patch is as good as sampling at all the patch's points – the mean is unlikely to differ (except at raster pixel boundaries).

On the other hand, a raster with a 10m resolution (e.g. Sentinel 1 and 2 data) may be worthwhile to sample at all the locations comprising a residence patch, so as to calculate the mean and variance of environmental conditions.

Furthermore, many (if not all) animals integrate cues from quite a distance (10m – 100m) when making decisions on when to settle in an area, and when to leave. Thus it can also be useful to sample environmental layers not at point locations, but to extract the mean and variance from an *area*, or a buffer, around the animal's point locations.

We have provided a convenient function to get either *(1)* the points (classified into patches), or *(2)* a summary ouput of the residence patches (i.e., the median coordinates and their attributes), or finally *(3)* a spatial buffer around the points from *(1)*. This function, `atl_patch_summary` implements these options using the `which_data` argument, where the options are *(1)* "points", *(2)* "summary", or *(3)* "spatial".

Here, we choose option (3), using a spatial buffer of 20m. The distance of the bufffer is passed to the argument `buffer_radius`.

```
# for the known and unkniwn patches
patch_sf_data <- atl_patch_summary(patch_res_known,
  which_data = "spatial",
  buffer_radius = 20
)

# assign crs
sf::st_crs(patch_sf_data) <- 32631

# get summary data
patch_summary_data <- atl_patch_summary(patch_res_known,
  which_data = "summary"
)
```

At this stage, users have successfully pre-processed their data from raw positions to residence patches. Residence patches are essentially `sf` objects and can be visualised using the `sf` method for `plot`; for instance `plot(patch_sf_data)`. Further sections reproduce the analyses in the main manuscript.

---

##### 1.9.6 Prepare to plot data

We read in the island's shapefile to plot it as a background for the residence patch figure.

```
# read griend and hut
griend <- sf::st_read("data/griend_polygon/griend_polygon.shp", quiet = TRUE)
hut <- sf::st_read("data/griend_hut.gpkg", quiet = TRUE)
```

#### 1.10 Compare patch metrics

We filter these data to exclude one exceedingly long outlier of about an hour (WP080).

```
# round median coordinate for inferred patches
patch_summary_inferred <-
  patch_summary_data[
    ,
    c(
      "x_median", "y_median",
      "nfixes", "duration", "patch"
    )
  ][, `:=`(
    x_median = round(x_median, digits = -2),
    y_median = round(y_median, digits = -2)
  )]
```

We add data from the known patches, matching by X and Y median.

```
# join with respatch summary
patch_summary_compare <-
  merge(patch_summary_real,
    patch_summary_inferred,
    on = c("x_median", "y_median"),
    all.x = TRUE, all.y = TRUE
  )

# drop nas
patch_summary_compare <- na.omit(patch_summary_compare)

# drop patch around WP080
patch_summary_compare <- patch_summary_compare[tID != "WP080", ]
```

7 patches are identified where there are no waypoints, while 2 waypoints are not identified as patches. These waypoints consisted of 6 and 15 (WP098 and WP092) positions respectively, and were lost when the data were aggregated to 30 second intervals.

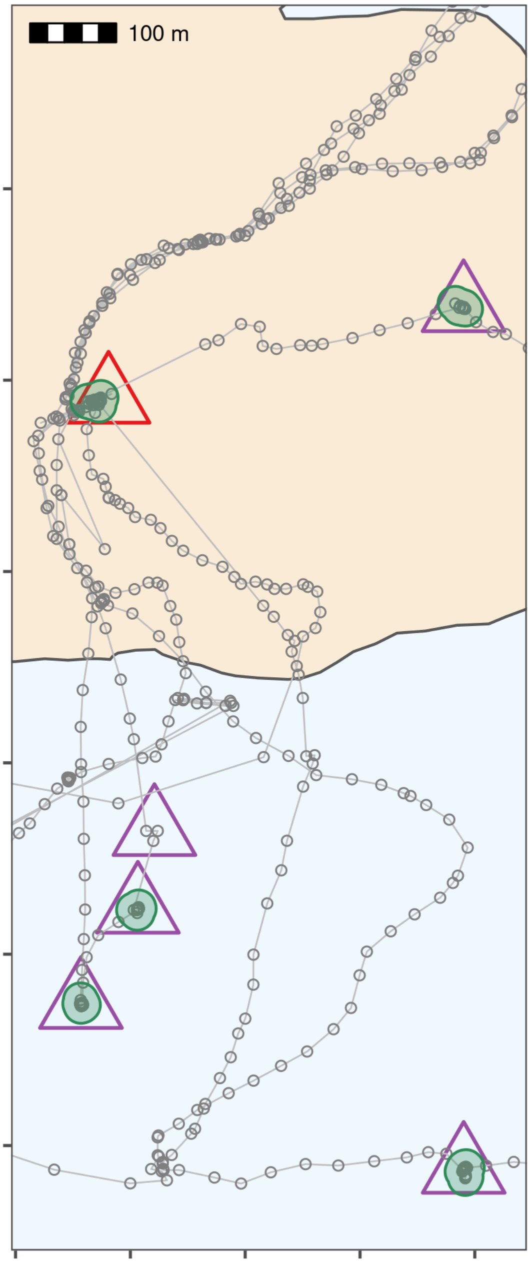

Figure 1.6: Classifying thinned data into residence patches yields robust estimates of the duration of known stops. The island of Griend (53.25°N, 5.25°E) is shown in beige. Residence patches (green polygons; function parameters in text) correspond well to the locations of known stops (purple triangles). However, the algorithm identified all areas with prolonged residence, including those which were not intended stops (n = 12; green polygons without triangles). The field station on Griend (red triangle) was not intended to be a stop, but the tags were stored here before the trial, and the method correctly picked up this prolonged stationary data as a residence patch. The algorithm failed to find two stops of 6 and 15 seconds duration, since these were lost in the data thinning step (purple triangle without green polygon shows one of these). The area shown is the lower rectangle in Fig. 1.3.

##### 219 1.10.1 Linear model durations

220 We run a simple linear model.

```
# get linear model
model_duration <- lm(duration_real ~ duration,
  data = patch_summary_compare
)

# get R2
summary(model_duration)
#>
#> Call:
#> lm(formula = duration_real ~ duration, data = patch_summary_compare)
#>
#> Residuals:
#>     Min      1Q  Median      3Q     Max
#> -105.07  -16.51   -4.26    9.54   91.66
#>
#> Coefficients:
#>             Estimate Std. Error t value Pr(>|t|)
#> (Intercept) 102.4395    47.4097    2.16    0.046 *
#> duration      1.0225     0.0786   13.02  6.3e-10 ***
#> ---
#> Signif. codes:  0 '***' 0.001 '**' 0.01 '*' 0.05 '.' 0.1 ' ' 1
#>
#> Residual standard error: 50.2 on 16 degrees of freedom
#> Multiple R-squared:  0.914,  Adjusted R-squared:  0.908
#> F-statistic:  169 on 1 and 16 DF,  p-value: 6.29e-10

# write to file
writeLines(
  text = capture.output(
    summary(model_duration)
  ),
  con = "data/model_output_residence_patch.txt"
)
```

##### 221 1.10.2 Linear model summary

```
cat(
  readLines(
    con = "data/model_output_residence_patch.txt",
    encoding = "UTF-8"
  ),
  sep = "\n"
)
#>
#> Call:
#> lm(formula = duration_real ~ duration, data = patch_summary_compare)
#>
#> Residuals:
```

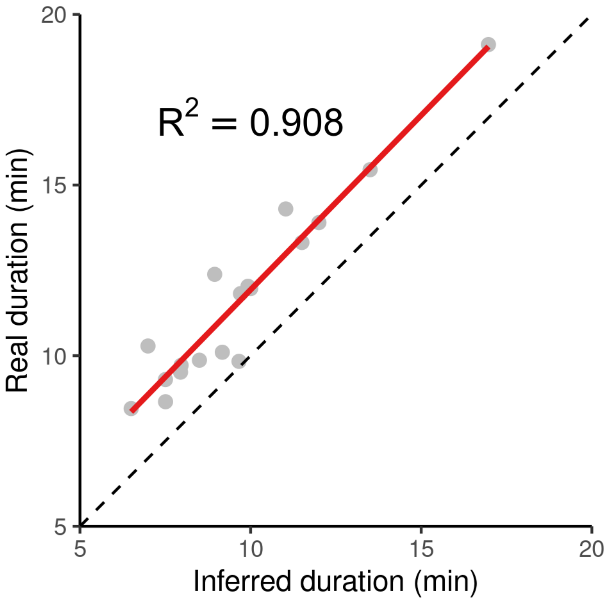

Figure 1.7: The inferred duration of residence patches corresponds very closely to the real duration (grey circles, red line shows linear model fit), with an underestimation of the true duration of around 2%. The dashed black line represents $y = x$ for reference.

```
#>     Min      1Q  Median      3Q     Max
#> -105.07  -16.51   -4.26    9.54   91.66
#>
#> Coefficients:
#>             Estimate Std. Error t value Pr(>|t|)
#> (Intercept) 102.4395    47.4097    2.16    0.046 *
#> duration      1.0225     0.0786   13.02  6.3e-10 ***
#> ---
#> Signif. codes:  0 '***' 0.001 '**' 0.01 '*' 0.05 '.' 0.1 ' ' 1
#>
#> Residual standard error: 50.2 on 16 degrees of freedom
#> Multiple R-squared:  0.914,  Adjusted R-squared:  0.908
#> F-statistic:  169 on 1 and 16 DF,  p-value: 6.29e-10
```

#### 1.11 Main text Figure 6

Plotting code is not shown in PDF and HTML form, see the `.Rmd` file.

### 2 Processing Egyptian Fruit Bat Tracks

We show the pre-processing pipeline at work on the tracks of three Egyptian fruit bats (*Rousettus aegyptiacus*), and construct residence patches.

#### 2.1 Prepare libraries

Install the required `R` libraries that are required from CRAN if not already installed.

```
# libs for data
library(data.table)
library(RSQLite)
library(atlastools)

# libs for plotting
library(ggplot2)
library(patchwork)

# recursion analysis
library(recurse)

# prepare a palette
pal <- RColorBrewer::brewer.pal(4, "Set1")
```

```
if (!require(remotes)) {
  install.packages("remotes", repos = "http://cran.us.r-project.org")
}

# installation using remotes
remotes::install_github("pratikunterwegs/atlastools")
```

#### 2.2 Read bat data

Read the bat data from an `SQLite` database local file and convert to a plain text csv file. This data can be found in the "data" folder.

```
# prepare the connection
con <- dbConnect(
  drv = SQLite(),
  dbname = "data/Three_example_bats.sql"
)

# list the tables
table_name <- dbListTables(con)
```

```
# prepare to query all tables
query <- sprintf('select * from \"%s\"', table_name)

# query the database
data <- dbGetQuery(conn = con, statement = query)

# disconnect from database
dbDisconnect(con)
```

Convert data to csv, and save a local copy in the folder “data”.

```
# convert data to datatable
setDT(data)

# write data for QGIS
fwrite(data, file = "data/bat_data.csv")
```

#### 2.3 Exploratory Data Analysis Panels: Main Text Figure 1

Here, we make some basic figures for exploratory data analysis shown in Figure 1 of the main text.

Plot the bat data as a sanity check, and inspect it visually for errors. The plot code is hidden in the rendered copy (PDF) of this supplementary material, but is available in the Rmarkdown file “supplement/06_bat_data.Rmd”.

##### 2.3.1 Heatmap of Locations

Here we demonstrate a basic heatmap of locations, aggregating over all individuals. In this one instance, the plotting code is also shown as a guide for readers, but in general, plotting code is hidden throughout this document.

```
data_heatmap <- copy(data)
data_heatmap[, c("xround", "yround") := list(
  plyr::round_any(X, 250),
  plyr::round_any(Y, 250)
)]
data_heatmap <- data_heatmap[, .N, by = c("xround", "yround")]

fig_heatmap <-
  ggplot(data_heatmap) +
  geom_tile(
    aes(
      xround, yround,
      fill = N
    ),
    colour = "grey",
    size = 0.1,
    show.legend = F
  ) +
  scale_fill_viridis_c(
    option = "C",
    direction = 1,
```

```
    trans = "log10"
  ) +
  ggthemes::theme_few() +
  theme(
    axis.text = element_blank(),
    axis.title = element_blank()
  ) +
  coord_sf(crs = 2039)

ggsave(fig_heatmap,
  filename = "supplement/figures/fig_bat_heatmap_raw.png",
  dpi = 300,
  width = 6, height = 4
)
```

##### 2.3.2 Sampling Intervals

Here, we create the histogram of sampling intervals shown in Figure 1 of the main text. The plotting code is hidden in the PDF version, but available in the source code.

##### 2.3.3 Localisation Error Measured by Systems

Here, we create the histogram of location error (variance in X) (Weiser et al. 2016) shown in Figure 1 of the main text. The plotting code is hidden in the PDF version, but available in the source code.

##### 2.3.4 Plot paths from raw tracking data

Here, we plot the paths of individual bats from the raw tracking data to visually inspect them for errors.

```
# this figure for the panel in main text figure 1
fig_bat_focus_bad_speed <-
  fig_bat_raw +
  coord_sf(
    crs = 2039,
    xlim = c(253000, NA),
    ylim = c(772000, NA)
  )

# save to supplement figures
ggsave(fig_bat_focus_bad_speed,
  filename = "supplement/figures/fig_bat_focus_bad_speed.png",
  dpi = 300,
  width = 4, height = 6
)
```

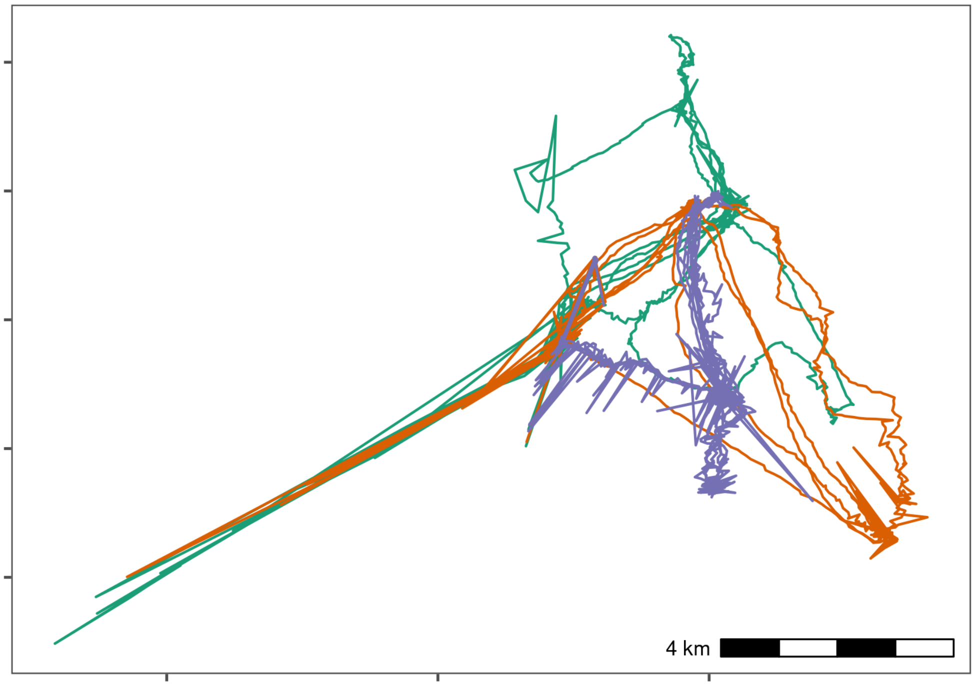

Figure 2.1: Movement data from three Egyptian fruit bats tracked using the ATLAS system (*Rousettus aegyptiacus*; (Toledo et al. 2020; Shohami and Nathan 2020)). The bats were tracked in the Hula Valley, Israel (33.1°N, 35.6°E), and we use three nights of tracking (5$^{th}$, 6$^{th}$, and 7$^{th}$ May, 2018), for our demonstration, with an average of 13,370 positions (SD = 2,173; range = 11,195 – 15,542; interval = 8 seconds) per individual. After first plotting the individual tracks, we notice severe distortions, making pre-processing necesary

#### 2.4 Prepare data for filtering

Here we apply a series of simple filters. It is always safer to deal with one individual at a time, so we split the data.table into a list of data.tables to avoid mixups among individuals.

This is a very rudimentary demonstration of the principle behind **batch processing** — splitting data into smaller, independent subsets, and applying the same steps to each subset.

##### 2.4.1 Prepare data per individual

```
# split bat data by tag
# first make a copy using the data.table function copy
# this prevents the orignal data from being modified by atlastools
# functions which DO MODIFY BY REFERENCE!
data_split <- copy(data)

# now split
data_split <- split(data_split, by = "TAG")
```

#### 2.5 Filter by system-generated error attributes

No natural bounds suggest themselves, so instead we proceed to filter by system-generated attributes of error, since point outliers are obviously visible.

We use filter out positions with `SD > 20` and positions calculated using only 3 base stations, using the function `atl_filter_covariates`.

First we calculate the variable `SD`, which for ATLAS systems is calculated as:

$$SD = \sqrt{VARX + VARY + 2 \times COVXY}$$

```
# get SD.
# since the data are data.tables, no assignment is necessary
invisible(
  lapply(data_split, function(dt) {
    dt[, SD := sqrt(VARX + VARY + (2 * COVXY))]
  })
)
```

Then we pass the filters to `atl_filter_covariates`. We apply the filter to each individual's data using an `lapply` – this separates the data from each individual into a separate data frame, lessening the chances of inter-individual mix-ups.

This is another basic example of the principles behind batch-processing, and could be parallelised using the `R` package `furrr` (see https://CRAN.R-project.org/package=furrr).

```
# filter for SD <= 20
# here, reassignment is necessary as rows are being removed
data_split <- lapply(data_split, function(dt) {
  dt <- atl_filter_covariates(
```

```
    data = dt,
    filters = c(
      "SD <= 20",
      "NBS > 3"
    )
  )
})
```

##### 2.5.1 Sanity check: Plot filtered data

We plot the data to check whether the filtering has improved the data (Fig. 2.2). The plot code is once again hidden in this rendering, but is available in the source code file.

#### 2.6 Filter by speed

Some point outliers remain, and could be removed using a speed filter.

First we calculate speeds, using `atl_get_speed`. We must assign the speed output to a new column in the data.table, which has a special syntax which modifies in place, and is shown below. This syntax is a feature of the `data.table` package, not strictly of `atlastools` (Dowle and Srinivasan 2020).

```
# get speeds as with SD, no reassignment required for columns
invisible(
  lapply(data_split, function(dt) {

    # first process time to seconds
    # assign to a new column
    dt[, time := floor(TIME / 1000)]

    dt[, `:=`(
      speed_in = atl_get_speed(dt,
        x = "X", y = "Y",
        time = "time",
        type = "in"
      ),
      speed_out = atl_get_speed(dt,
        x = "X", y = "Y",
        time = "time",
        type = "out"
      )
    )]
  })
)
```

Now filter for speeds > 20 m/s (around 70 km/h), passing the predicate (a statement return TRUE or FALSE) to `atl_filter_covariates`. First, we remove positions which have `NA` for their `speed_in` (the first position) and their `speed_out` (last position).

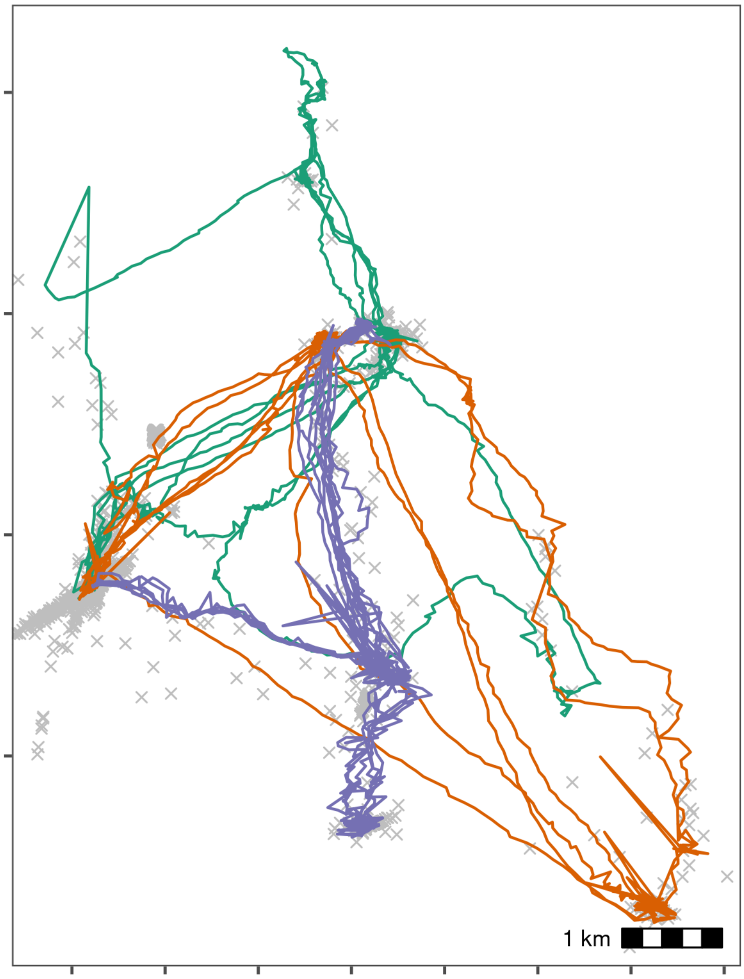

Figure 2.2: Bat data filtered for large location errors, removing observations with standard deviation > 20. Grey crosses show data that were removed. Since the number of base stations used in the location process is a good indicator of error (Weiser et al. 2016), we also removed observations calculated using fewer than four base stations. Both steps used the function `atl_filter_covariates`. This filtering reduced the data to an average of 10,447 positions per individual (78% of the raw data on average). However, some point outliers remain.

```r
# filter speeds
# reassignment is required here
data_split <- lapply(data_split, function(dt) {
  dt <- na.omit(dt, cols = c("speed_in", "speed_out"))

  dt <- atl_filter_covariates(
    data = dt,
    filters = c(
      "speed_in <= 20",
      "speed_out <= 20"
    )
  )
})
```

##### 2.6.1 Sanity check: Plot speed filtered data

The speed filtered data is now inspected for errors (Fig. 2.3). The plot code is once again hidden.

#### 2.7 Median smoothing

The quality of the data is relatively high, and a median smooth is not strictly necessary. We demonstrate the application of a 5 point median smooth to the data nonetheless (Fig. 2.4).

Since the median smoothing function `atl_median_smooth` modifies in place, we first make a copy of the data, using `data.table`'s `copy` function. No reassignment is required, in this case. The `lapply` function allows arguments to `atl_median_smooth` to be passed within `lapply` itself.

In this case, the same moving window $K$ is applied to all individuals, but modifying this code to use the multivariate version `Map` allows different $K$ to be used for different individuals. This is a programming matter, and is not covered here further.

```r
# since the function modifies in place, we shall make a copy
data_smooth <- copy(data_split)

# split the data again
data_smooth <- split(data_smooth, by = "TAG")

# apply the median smooth to each list element
# no reassignment is required as THE FUNCTION MODIFIES IN PLACE!
invisible(

  # the function arguments to atl_median_smooth
  # can be passed directly in lapply

  lapply(
    X = data_smooth,
    FUN = atl_median_smooth,
    time = "time", moving_window = 5
```

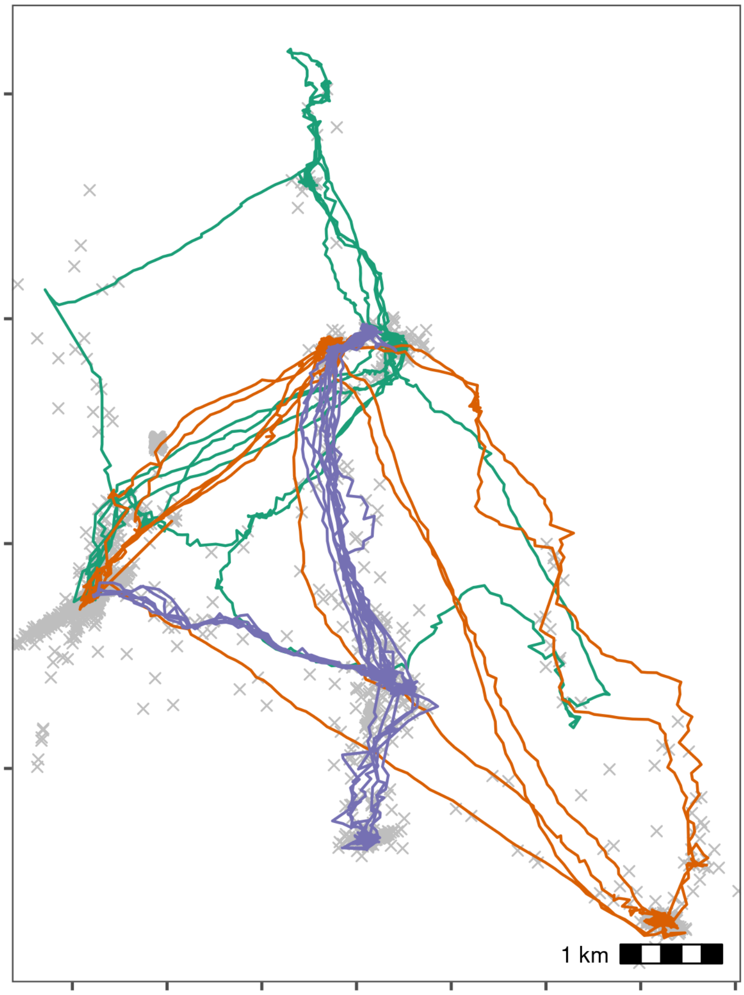

Figure 2.3: Bat data with unrealistic speeds removed. Notice, compared with the previous figure, that spikes of unrealistic movement in all three tracks have been removed. Grey crosses show data that were removed. We calculated the incoming and outgoing speed of each position using `atl_get_speed`, and filtered out positions with speeds > 20 m/s using `atl_filter_covariates`, leaving 10,337 positions per individual on average (98% from the previous step).

```
  )
)
```

##### 2.7.1 Sanity check: Plot smoothed data

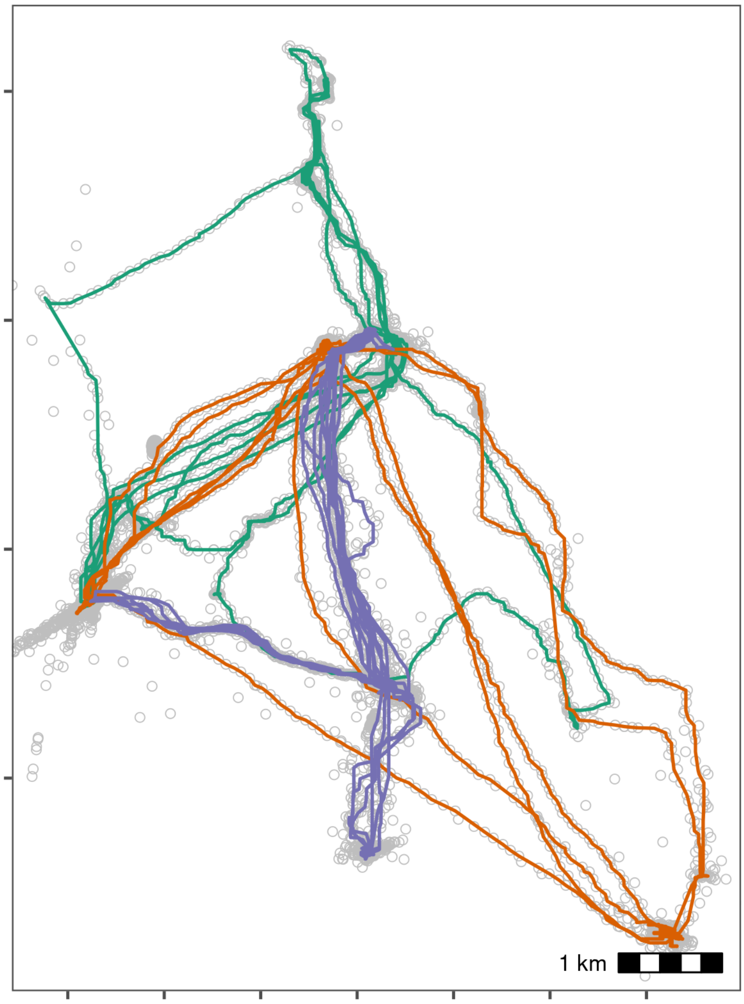

Figure 2.4: Bat data after applying a median smooth with a moving window $K = 5$. Grey circles show data prior to smoothing. The smoothing step did not discard any data.

#### 2.8 Making residence patches

##### 2.8.1 Calculating residence time

First, the data is put through the `recurse` package to get residence time (Bracis, Bildstein, and Mueller 2018).

```
# split the data
data_smooth <- split(data_smooth, data_smooth$TAG)
```

We calculated residence time, but since bats may revisit the same features, we want to prevent confusion between frequent revisits and prolonged residence.

For this, we stop summing residence times within $Z$ metres of a location if the animal exited the area for one hour or more. The value of $Z$ (radius, in `recurse` parameter terms) was chosen as 50m.

This step is relatively complicated and is only required for individuals which frequently return to the same location, or pass over the same areas repeatedly, and for which revisits (cumulative time spent) may be confused for residence time in a single visit.

While a simpler implementation using total residence time divided by the number of revisits is also possible, this does assume that each revisit had the same residence time.

```
# get residence times
data_residence <- lapply(data_smooth, function(dt) {

  # do basic recurse -- refer to Bracis et al. (2018) Ecography
  dt_recurse <- getRecursions(
    x = dt[, c("X", "Y", "time", "TAG")],
    radius = 50,
    timeunits = "mins"
  )

  # get revisit stats column provided as recurse output
  dt_recurse <- setDT(
    dt_recurse[["revisitStats"]]
  )

  # count long absences from the each position
  dt_recurse[, timeSinceLastVisit :=
    ifelse(is.na(timeSinceLastVisit), -Inf, timeSinceLastVisit)]
  dt_recurse[, longAbsenceCounter := cumsum(timeSinceLastVisit > 5),
    by = .(coordIdx)
  ]

  # filter data to exclude revisits after the first long absence of 60 mins
  dt_recurse <- dt_recurse[longAbsenceCounter < 1, ]

  # calculate the residence time as the sum of times inside
  # before the first 'long absence'
  # also calculate the First Passage Time and the number of revisits
  dt_recurse <- dt_recurse[, list(
    resTime = sum(timeInside),
    fpt = first(timeInside),
    revisits = max(visitIdx)
  ),
  by = .(coordIdx, x, y)
  ]

  # prepare to merge existing data with recursion data
```

```
  dt[, coordIdx := seq(nrow(dt))]

  # merge the revised recursion analysis data with the tracking data
  dt <- merge(dt,
    dt_recurse[, c("coordIdx", "resTime")],
    by = c("coordIdx")
  )

  # ensure the data are ordered in ascending order of time
  setorderv(dt, "time")

  # print message when done
  message(sprintf("TAG %s residence times done", unique(dt$TAG)))

  # return the dataframe
  dt
})
```

We bind the data together and assign a human readable timestamp column.

```
# bind the list
data_residence <- rbindlist(data_residence)

# get time as human readable
data_residence[, ts := as.POSIXct(time, origin = "1970-01-01")]

# get hour of day to filter for nighttime
data_residence[, hour := data.table::hour(ts)]

# filter for hour between 8pm and 5am
data_residence <- atl_filter_covariates(
  data = data_residence,
  filters = "hour > 20 | hour < 5"
)
```

##### 2.8.2 Constructing residence patches

Some preparation is required. First, the function requires columns `x`, `y`, `time`, and `id`, which we assign using the `data.table` syntax. The `time` column is already present, but the other columns need to be renamed to lower case. Then we subset the data to only work with positions where the individual had a residence time of more than 5 minutes.

```
# add an id column
data_residence[, `:=`(
  id = TAG,
  x = X, y = Y
)]

# get mean residence time per id
data_residence[, list(
  mean_residence = mean(resTime),
  sd_residence = sd(resTime)
```

```
), by = "TAG"]
#>             TAG mean_residence sd_residence
#> 1: 972001004424          202.1        135.8
#> 2: 972001004449           28.1         34.0
#> 3: 972001004452           43.5         47.3

# get mean residence time for all bats pooled
data_residence[, list(
  mean_residence = mean(resTime),
  sd_residence = sd(resTime)
)]
#>    mean_residence sd_residence
#> 1:           95.6          119

# filter for residence time > 5 minutes
data_residence <- data_residence[resTime > 5, ]

# average positions per bat after removing transit points
data_residence[, list(.N), by = "TAG"][, list(mean_positions_ = mean(N))]
#>    mean_positions_
#> 1:            5736

# split the data
data_residence <- split(data_residence, data_residence$TAG)
```

We apply the residence patch method, using the default argument values (`lim_spat_indep = 100` (metres), `lim_time_indep = 30` (minutes)). We change the `buffer_radius` to 25 metres (twice the buffer radius is used, so points must be separated by 50m to be independent bouts), and `min_fixes = 3`.

```
# segment into residence patches
data_patches <- lapply(data_residence, atl_res_patch,
  buffer_radius = 25,
  min_fixes = 3
)
```

##### 2.8.3 Getting residence patch data

We extract the residence patch data as spatial `sf-MULTIPOLYGON` objects. These are returned as a list and must be converted into a single `sf` object. These objects and the raw movement data are shown in Fig. 2.5.

```
# get data spatials
data_spatials <- lapply(data_patches, atl_patch_summary,
  which_data = "spatial",
  buffer_radius = 25
)

# bind list
data_spatials <- rbindlist(data_spatials)

# convert to sf
library(sf)
```

```r
data_spatials <- st_sf(data_spatials, sf_column_name = "polygons")

# assign a crs
st_crs(data_spatials) <- st_crs(2039)
```

##### 2.8.4 Write patch spatial representations

```r
st_write(data_spatials,
  dsn = "data/data_bat_residence_patches.gpkg",
  append = FALSE
)
#> Deleting layer `data_bat_residence_patches' using driver `GPKG'
#> Writing layer `data_bat_residence_patches' to data source `data/data_bat_residence_pa
#> Writing 55 features with 18 fields and geometry type Multi Polygon.
```

Write cleaned bat data.

```r
fwrite(rbindlist(data_smooth),
  file = "data/data_bat_smooth.csv"
)
```

Write patch summary.

```r
# get summary
patch_summary <- lapply(data_patches, atl_patch_summary)

# bind summary
patch_summary <- rbindlist(patch_summary)

# write
fwrite(
  patch_summary,
  "data/data_bat_patch_summary.csv"
)
```

##### 2.8.5 Duration at foraging sites

We exclude the first and last patch of each day as being roosting related, and examine how much of the total foraging time (time between the remaining first and last patch) was spent at foraging sites. It follows that the remainder of the time must have been spent in transit, or otherwise not foraging.

```r
# make patch summary a datatable
setDT(patch_summary)

# exclude roost
patch_summary <- patch_summary[x_median > 254000, ]

# get mean and sd of duration in patches
patch_summary[, list(
  mean_duration = mean(duration / 60),
  sd_duration = sd(duration / 60)
)]
```

```r
#>    mean_duration sd_duration
#> 1:            57        62.2

# assign night id
patch_summary[, c("hour", "day") := list(
  data.table::hour(as.POSIXct(time_start, origin = "1970-01-01")),
  data.table::mday(as.POSIXct(time_start, origin = "1970-01-01"))
)]

patch_summary[, night := 1 + c(0, cumsum(diff(hour) > 12)), by = "id"]

# get total foraging time
foraging_proportion <- patch_summary[, list(
  time_total_forage = (max(time_end) - min(time_start)) / 60,
  time_forage_site = sum(duration / 60)
),
by = c("id", "night")
]

# get proporion of foraging that is at a foraging site
foraging_proportion[, list(
  mean_foraging_prop = mean(time_forage_site / time_total_forage),
  sd_foraging_prop = sd(time_forage_site / time_total_forage)
)]
#>    mean_foraging_prop sd_foraging_prop
#> 1:              0.838            0.155
```

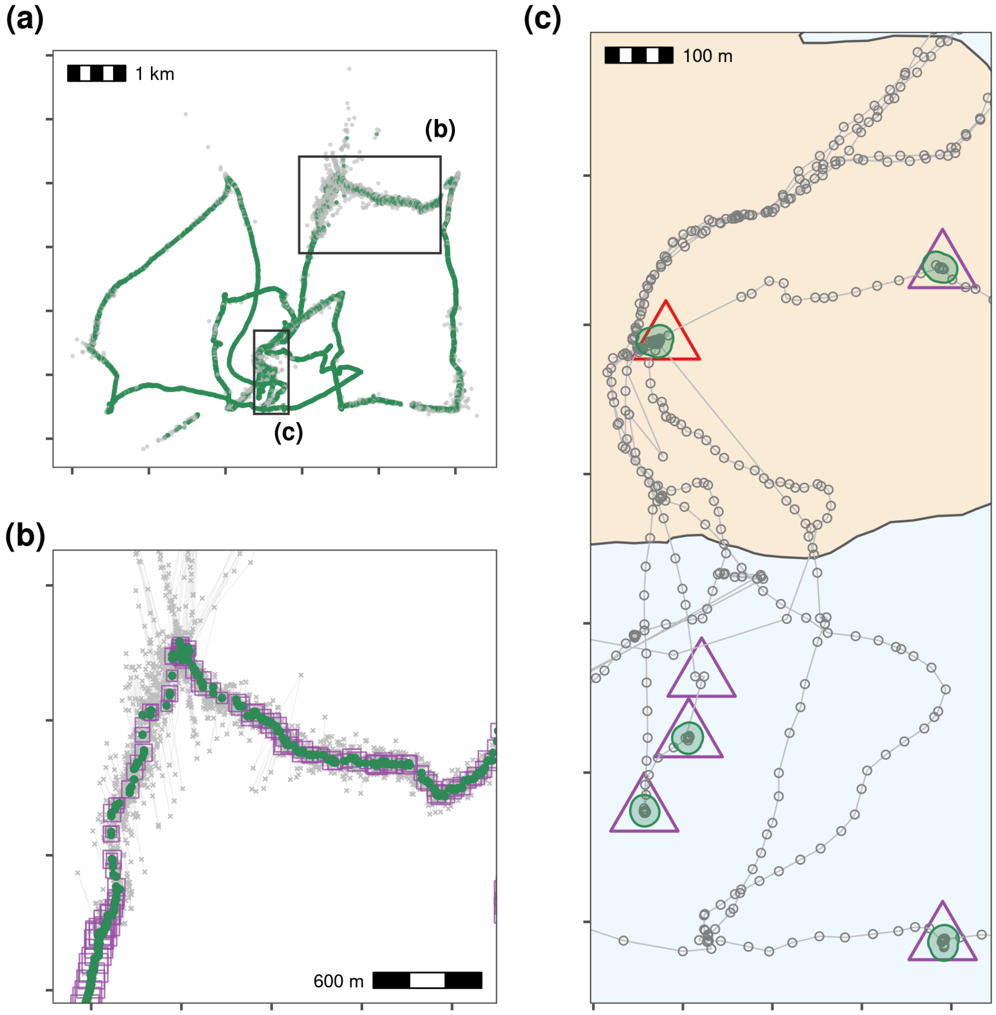

Figure 2.5: A visual examination of plots of the bats' residence patches and linear approximations of paths between them showed that though all three bats roosted at the same site, they used distinct areas of the study site over the three nights **(a)**. Bats tended to be resident near fruit trees, which are their main food source, travelling repeatedly between previously visited areas **(b, c)**. However, bats also appeared to spend some time at locations where no fruit trees were recorded, prompting questions about their use of other food sources **(b, c)**. When bats did occur close together, their residence patches barely overlapped, and their paths to and from the broad area of co-occurrence were not similar **(c)**. Constructing residence patches for multiple individuals over multiple activity periods suggests interesting dynamics of within- and between-individual overlap **(b, c)**.

Dowle, Matt, and Arun Srinivasan. 2020. *Data.Table: Extension of 'data.Frame'*. Manual.

Gupte, Pratik Rajan. 2020. "Atlastools: Pre-Processing Tools for High Frequency Tracking Data." Zenodo. https://doi.org/10.5281/ZENODO.4033154.

Oudman, Thomas, Theunis Piersma, Mohamed V. Ahmedou Salem, Marieke E. Feis, Anne Dekinga, Sander Holthuijsen, Job ten Horn, Jan A. van Gils, and Allert I. Bijleveld. 2018. "Resource Landscapes Explain Contrasting Patterns of Aggregation and Site Fidelity by Red Knots at Two Wintering Sites." *Movement Ecology* 6 (1): 24–24. https://doi.org/10.1186/s40462-018-0142-4.

Shohami, David, and Ran Nathan. 2020. "Cognitive Map-Based Navigation in Wild Bats Revealed by a New High-Throughput Tracking System." Dryad. https://doi.org/10.5061/DRYAD.G4F4QRFN2.

Toledo, Sivan, David Shohami, Ingo Schiffner, Emmanuel Lourie, Yotam Orchan, Yoav Bartan, and Ran Nathan. 2020. "Cognitive MapBased Navigation in Wild Bats Revealed by a New High-Throughput Tracking System." *Science* 369 (6500). American Association for the Advancement of Science: 188–93. https://doi.org/10.1126/science.aax6904.

Weiser, Adi Weller, Yotam Orchan, Ran Nathan, Motti Charter, Anthony J. Weiss, and Sivan Toledo. 2016. "Characterizing the Accuracy of a Self-Synchronized Reverse-GPS Wildlife Localization System." In *2016 15th ACM/IEEE International Conference on Information Processing in Sensor Networks (IPSN)*, 1–12. https://doi.org/10.1109/IPSN.2016.7460662.
