## Supplementary Material 2 for "A Guide to Pre-Processing High-Throughput Animal Tracking Data"

### Package ‘atlastools’

July 1, 2021

**Title** Tools for Pre-Processing High-Throughput Animal Tracking Data

**Version** 1.0.0000

**Description** Tools to pre-process high-throughput tracking data. Developed for use with ATLAS tracking systems, but suitable for use with nearly any XY-time animal tracking data.

**License** GPL-3

**Encoding** UTF-8

**LazyData** true

**URL** https://github.com/pratikunterwegs/atlastools

**BugReports** https://github.com/pratikunterwegs/atlastools/issues

**Imports** sf,
glue,
data.table,
stats,
assertthat,
stringr,
bit64

**RoxygenNote** 7.1.1

**Roxygen** list(markdown = TRUE)

**Suggests** testthat (>= 2.1.0),
ggplot2,
scales,
knitr,
rmarkdown

**VignetteBuilder** knitr

#### R topics documented:

**Index** **16**

---

`atl_check_data` *Check data has required columns.*

---

#### Description

An internal function that checks that the data.table has the required columns. Used within most, if not all, other atlastools functions.

#### Usage

```
atl_check_data(data, names_expected = c("x", "y", "time"))
```

#### Arguments

| | |
|---|---|
| `data` | The tracking data to check for required columns. Must be in the form of a data.frame or similar, which can be handled by the function colnames. |
| `names_expected` | The names expected as a character vector. By default, checks for the column names `x,y,time`. |

#### Value

None. Breaks if the data does not have required columns.

#### Author(s)

Pratik R. Gupte

#### Examples

```
# basic (and only) use
## Not run:
atl_check_data(
  data = data,
  names_expected = c("x", "y", "time")
)

## End(Not run)
```

atl_filter_bounds *Filter positions by an area.*

#### Description

Filters out positions lying inside or outside an area. The area can be defined in two ways, either by its X and Y coordinate ranges, or by an `sf-*POLYGON` object. `MULTIPOLYGON` objects are supported by the internal function `atl_within_polygon`.

#### Usage

```
atl_filter_bounds(
  data,
  x = "x",
  y = "y",
  x_range = NA,
  y_range = NA,
  sf_polygon = NULL,
  remove_inside = TRUE
)
```

#### Arguments

| | |
|---|---|
| data | A dataframe or extension which contains X and Y coordinates. |
| x | The X coordinate column. |
| y | The Y coordinate column. |
| x_range | The range of X coordinates. |
| y_range | The range of Y coordinates. |
| sf_polygon | `sfc_*POLYGON` object which must have a defined CRS. The polygon CRS is assumed to be appropriate for the positions as well, and is assigned to the coordinates when determining the intersection. |
| remove_inside | Whether to remove points from within the range. Setting `negate = TRUE` removes positions within the bounding box specified by the X and Y ranges. |

#### Value

A data frame of tracking locations with attractor points removed.

#### Author(s)

Pratik R. Gupte

#### Examples

```
## Not run:
filtered_data <- atl_filter_bounds(
  data = data,
  x = "X", y = "Y",
  x_range = c(x_min, x_max),
  y_range = c(y_min, y_max),
  sf_polygon = your_polygon,
  remove_inside = FALSE
)

## End(Not run)
```

---

`atl_filter_covariates` *Filter data by position covariates.*

---

#### Description

The atlastools function `atl_filter_covariates` allows convenient filtering of a dataset by any number of logical filters. This function can be used to easily filter timestamps in a range, as well as combine simple spatial and temporal filters. It accepts a character vector of `R` expressions that each return a logical vector (i.e., `TRUE` or `FALSE`). Each filtering condition is interpreted in the context of the dataset supplied, and used to filter for rows that satisfy each of the filter conditions. Users must make sure that the filtering variables exist in their dataset in order to avoid errors.

#### Usage

```
atl_filter_covariates(data, filters = c())
```

#### Arguments

| | |
|---|---|
| `data` | A dataframe or similar containing the variables to be filtered. |
| `filters` | A character vector of filter expressions. An example might be `"speed < 20"`. The filtering variables must be in the dataframe. The function will not explicitly check whether the filtering variables are present; this makes it flexible, allowing expressions such as `"between(speed,2,20)"`, but also something to use at your own risk. A missing filter variables *will* result in an empty data frame. |

#### Value

A dataframe filtered using the filters specified.

#### Author(s)

Pratik R. Gupte

#### Examples

```
## Not run:
night_data <- atl_filter_covariates(
  data = dataset,
  filters = c("!inrange(hour, 6, 18)")
)

data_in_area <- atl_filter_covariates(
  data = dataset,
  filters = c(
    "between(time, t_min, t_max)",
    "between(x, x_min, x_max)"
  )
)
filtered_data <- atl_filter_covariates(
  data = data,
  filters = c(
    "NBS > 3",
    "SD < 100",
    "between(day, 5, 8)"
  )
)

## End(Not run)
```

---

`atl_get_speed` *Calculate instantaenous speed.*

---

#### Description

Returns speed in metres per time interval. The time interval is dependent on the units of the column specified in `time`. Users should apply this function to *one individual at a time*, ideally by splittng a dataframe with multiple individuals into a list of dataframes.

#### Usage

```
atl_get_speed(data, x = "x", y = "y", time = "time", type = c("in"))
```

#### Arguments

| | |
|---|---|
| `data` | A dataframe or similar which must have the columns specified by `x`, `y`, and `time`. |
| `x` | The x coordinate. |
| `y` | The y coordinate. |
| `time` | The timestamp in seconds since the UNIX epoch. |
| `type` | The type of speed (incoming or outgoing) to return. Incoming speeds are specified by `type = "in"`, and outgoing speeds by `type = "out"`. |

#### Value

A vector of numerics representing speed. The first position is assigned a speed of NA.

#### Author(s)

Pratik R. Gupte

#### Examples

```
## Not run:
data$speed_in <- atl_get_speed(data,
  x = "x", y = "y",
  time = "time", type = c("in")
)

## End(Not run)
```

---

`atl_median_smooth` *Apply a median smooth to coordinates.*

---

#### Description

Applies a median smooth defined by a rolling window to the X and Y coordinates of the data. This function *modifies in place*, i.e., *the results need not be assigned to a new data.table*.

#### Usage

```
atl_median_smooth(data, x = "X", y = "Y", time = "TIME", moving_window = 3)
```

#### Arguments

| | |
|---|---|
| `data` | A dataframe object returned by getData. Must contain the columns "X", "Y", "SD", "NBS", "TAG", "TIME"; these are the X coordinate, Y coordinate, standard deviation in measurement, number of ATLAS towers that received the signal, the tag number, and the numeric time, in milliseconds from 1970-01-01. |
| `x` | The X coordinate. |
| `y` | The Y coordinate. |
| `time` | The timestamp, ideally as an integer. median calculation. |
| `moving_window` | The size of the moving window for the median smooth. Must be an odd number. |

#### Value

A datatable class object (extends data.frame) which has the additional columns posID and ts, which is TIME converted to human readable POSIXct format.

##### Examples

```
## Not run:
atl_median_smooth(
  data = track_data,
  x = "x", y = "y",
  time = "time",
  moving_window = 5
)

## End(Not run)
```

---

`atl_patch_dist` *Get the distance between patches.*

---

##### Description

Gets the linear distance between the first point of patch `i` and the last point of the previous patch patch `i -1`. Distance is returned in metres. This function is used internally by other functions, and rarely on its own.

##### Usage

```
atl_patch_dist(
  data,
  x1 = "x_end",
  x2 = "x_start",
  y1 = "y_end",
  y2 = "y_start"
)
```

##### Arguments

| | |
|---|---|
| `data` | A dataframe of or extending the class data.frame, such as a data.table. This must contain two pairs of coordinates, the start and end X and Y coordinates of a feature. |
| `x1` | The first X coordinate or longitude; for inter-patch distances, this is the last coordinate (x_end) of a patch $i$. |
| `x2` | The second X coordinate; for inter-patch distances, this is the first coordinate (x_start) of a subsequent patch $i + 1$. |
| `y1` | The first Y coordinate or latitude; for inter-patch distances, this is the last coordinate (y_end) of a patch $i$. |
| `y2` | The second Y coordinate; for inter-patch distances, this is the first coordinate (y_start) of a subsequent patch $i + 1$. |

#### Value

A numeric vector of the length of the number of patches, or rows in the input dataframe. For single patches, returns NA. The vector has as its elements NA, followed by n-1 distances, where n is the number of rows.

#### Author(s)

Pratik R. Gupte

#### Examples

```
# basic usage of atl_patch_dist
## Not run:
atl_patch_dist(
  data = data,
  x1 = "x_end", x2 = "x_start",
  y1 = "y_end", y2 = "y_start"
)

## End(Not run)
```

---

### atl_patch_summary *Get residence patch data.*

---

#### Description

The function atl_patch_summary can be used to extract patch-specific summary data such as the median coordinates, the patch duration, the distance travelled within the patch, the displacement within the patch, and the patch area.

#### Usage

```
atl_patch_summary(patch_data, which_data = "summary", buffer_radius = 10)
```

#### Arguments

patch_data
: A data.frame with a nested list column of the raw data underlying each patch. Since data.frames don't support nested columns, will actually be a data.table or similar extension.

which_data
: Which data to return. May be the raw data underlying the patch (which_data = "points"), or a spatial features (sf-MULTIPOLYGON) object with patch covariates (which_data = "spatial"), or a data.table of the patch covariates without the geometry column (which_data = "summary").

buffer_radius
: Spatial buffer radius (in metres) around points when requesting sf based polygons.

##### Value

An object of type `sf` or `data.table` depending on which data is requested.

##### Author(s)

Pratik R. Gupte

##### Examples

```
## Not run:
patch_summary <- atl_patch_summary(
  patch_data = patches,
  which_data = "summary",
  buffer_radius = 10
)

## End(Not run)
```

---

`atl_remove_reflections`    *Remove reflected positions.*

---

##### Description

Remove reflections, or prolonged spikes from a movement track by identifying the bounds and removing positions between them. The important function arguments here are `point_angle_cutoff` ($A$), `reflection_speed_cutoff` ($S$). If the prolonged spike ends before the last row of data, the true end point is used as the outer bound of the spike. If the prolonged spike does not end within the last row of data, all the data are retained and a message is printed.

##### Usage

```
atl_remove_reflections(
  data,
  x = "x",
  y = "y",
  time = "time",
  point_angle_cutoff = 45,
  reflection_speed_cutoff = 20
)
```

##### Arguments

| | |
|---|---|
| `data` | A dataframe or similar which has previously been cleaned. |
| `x` | The name of the X coordinate column. |
| `y` | The name of the Y coordinate column. |

`time` The name of the timestamp column.

`point_angle_cutoff`
The turning angle (in degrees) above which high instantaneous speeds are considered an anomaly rather than fast transit.

`reflection_speed_cutoff`
The speed (in m/s) above which an anomaly is detected when combined with a high turning angle.

#### Value

A dataframe with reflections removed.

#### Author(s)

Pratik R. Gupte

#### Examples

```
## Not run:
filtered_data <- atl_remove_reflections(
  data = track_data,
  x = "x", y = "y", time = "time",
  point_angle_cutoff = A,
  reflection_speed_cutoff = S
)

## End(Not run)
```

---

`atl_res_patch` *Construct residence patches from position data.*

---

#### Description

A cleaned movement track can be classified into residence patches using the function `atl_res_patch`. The function expects a specific organisation of the data: there should be at least the following columns, `x`, `y`, `time`, and `id`, all named in lower case, and corresponding to the coordinates, timestamp in the UNIX format (seconds since 1970), and the identity of the tracked individual. The result contains only the data that was classified as a residence patch and removes transit between them. `atl_res_patch` requires only three parameters: (1) the distance threshold between positions (called `buffer_size`), (2) the distance threshold between clusters of positions (called `lim_spat_indep`), and (3) the time interval between clusters (called `lim_time_indep`). Clusters formed of fewer than a minimum number of positions can be excluded. The exclusion of clusters with few positions can help in removing bias due to short stops, but if such short stops are also of interest, they can be included by reducing the `min_fixes` argument. Position covariates such as speed may also be summarised patch-wise by passing covariate names and summary functions as character vectors to the `summary_variables` and `summary_functions` arguments, respectively .

##### Usage

```
atl_res_patch(
  data,
  buffer_radius = 10,
  lim_spat_indep = 100,
  lim_time_indep = 30,
  min_fixes = 3,
  summary_variables = c(),
  summary_functions = c()
)
```

##### Arguments

| | |
|---|---|
| `data` | A dataframe of values of any class that is or extends data.frame. The dataframe must contain at least two spatial coordinates, `x` and `y`, and a temporal coordinate, `time`. The names of columns specifying these can be passed as arguments below. The column `id` indicating animal id is *required*. |
| `buffer_radius` | A numeric value specifying the radius of the buffer to be considered around each coordinate point. May be thought of as the distance that an individual can access, assess, or otherwise cover when at a discrete point in space. |
| `lim_spat_indep` | A numeric value of distance in metres of the spatial distance between two patches for them to the considered independent. |
| `lim_time_indep` | A numeric value of time in minutes of the time difference between two patches for them to be considered independent. |
| `min_fixes` | The minimum number of fixes for a group of spatially-proximate number of ponts to be considered a preliminary residence patch. |
| `summary_variables` | Optional variables for which patch-wise summary values are required. To be passed as a character vector. |
| `summary_functions` | The functions with which to summarise the summary variables; must return only a single value, such as median, mean etc. To be passed as a character vector. |

##### Value

A data.frame extension object. This dataframe has the added column `patch` and `patchdata`, indicating the patch identity and the data used to construct the patch. In addition, there are columns with patch summary variables.

##### Author(s)

Pratik R. Gupte

##### Examples

```
## Not run:
patches <- atl_res_patch(
```

```
  data = track_data,
  buffer_radius = 10,
  lim_spat_indep = 100,
  lim_time_indep = 30,
  min_fixes = 3,
  summary_variables = c("speed"),
  summary_functions = c("mean", "sd")
)

## End(Not run)
```

---

atl_simple_dist    *Calculate distances between successive points.*

---

#### Description

Gets the euclidean distance between consecutive points in a coordinate reference system in metres, i.e., UTM systems.

#### Usage

```
atl_simple_dist(data, x = "x", y = "y")
```

#### Arguments

| | |
|---|---|
| data | A dataframe object of or extending the class data.frame, which must contain two coordinate columns for the X and Y coordinates. |
| x | A column name in a data.frame object that contains the numeric X or longitude coordinate for position data. |
| y | A column name in a data.frame object that contains the numeric Y or latitude coordinate for position data. |

#### Value

Returns a vector of distances between consecutive points.

`atl_thin_data` *Thin tracking data by resampling or aggregation.*

#### Description

Uniformly reduce data volumes with either aggregation or resampling (specified by the `method` argument) over an interval specified in seconds using the `interval` argument. Both options make two important assumptions: (1) that timestamps are named time', and (2) all columns except the identity columns can be averaged in \code{R}. While the subsample' option returns a thinned dataset with all columns from the input data, the aggregate' option drops the column \code{COVXY}, since this cannot be propagated to the averaged position. Both options handle the column time' differently: while subsample' returns the actual timestamp (in UNIX time) of each sample, aggregate' returns the mean timestamp (also in UNIX time). In both cases, an extra column, `time_agg`, is added which has a uniform difference between each element corresponding to the user-defined thinning interval. The 'aggregate' option only recognises errors named `VARX` and `VARY`, and standard deviation around each position named `SD`. If all of these columns are not present together the function assumes there is no measure of error, and drops those columns. If there is actually no measure of error, the function simply returns the averaged position and covariates in each time interval. Grouping variables' names (such as animal identity) may be passed as a character vector to the `id_columns` argument.

#### Usage

```
atl_thin_data(
  data,
  interval = 60,
  id_columns = NULL,
  method = c("subsample", "aggregate")
)
```

#### Arguments

| | |
|---|---|
| `data` | Cleaned data to aggregate. Must have a numeric column named time. |
| `interval` | The interval in seconds over which to aggregate. |
| `id_columns` | Column names for grouping columns. |
| `method` | Should the data be thinned by subsampling or aggregation. If resampling (`method = "subsample"`), the first position of each group is taken. If aggregation (`method = "aggregate"`), the group positions' mean is taken. |

#### Value

A dataframe aggregated taking the mean over the interval.

#### Examples

```
## Not run:
thinned_data <- atl_thin_data(data,
  interval = 60,
```

```
  id_columns = c("animal_id"),
  method = "aggregate"
)

## End(Not run)
```

---

`atl_turning_angle` *Get the turning angle between points.*

---

#### Description

Gets the relative heading between two positions using the law of cosines. The turning angle is returned in degrees. Users should apply this function to *one individual at a time*, ideally by splittng a dataframe with multiple individuals into a list of dataframes.

#### Usage

```
atl_turning_angle(data, x = "x", y = "y", time = "time")
```

#### Arguments

| | |
|---|---|
| `data` | A dataframe or similar which must have the columns specified by `x`, `y`, and `time`. |
| `x` | The x coordinate. |
| `y` | The y coordinate. |
| `time` | The timestamp in seconds since the UNIX epoch. |

#### Value

A vector of turning angles in degrees. Negative degrees indicate 'left' turns. There are two fewer angles than the number of rows in the dataframe.

#### Author(s)

Pratik R. Gupte

#### Examples

```
## Not run:
data$angle <- atl_turning_angle(data,
  x = "x", y = "y", time = "time"
)

## End(Not run)
```

atl_within_polygon *Detect position intersections with a polygon.*

#### Description

Detects which positions intersect a `sfc_*POLYGON`. Tested only for single polygon objects.

#### Usage

```
atl_within_polygon(data, x = "x", y = "y", polygon)
```

#### Arguments

| | |
|---|---|
| `data` | A dataframe or similar containg at least X and Y coordinates. |
| `x` | The name of the X coordinate, assumed by default to be "x". |
| `y` | The Y coordinate as above, default "y". |
| `polygon` | An `sfc_*POLYGON` object which must have a defined CRS. The polygon CRS is assumed to be appropriate for the positions as well, and is assigned to the coordinates when determining the intersection. |

#### Value

Row numbers of positions which are inside the polygon.

### Index

atl_check_data, 2
atl_filter_bounds, 3
atl_filter_covariates, 4
atl_get_speed, 5
atl_median_smooth, 6
atl_patch_dist, 7
atl_patch_summary, 8
atl_remove_reflections, 9
atl_res_patch, 10
atl_simple_dist, 12
atl_thin_data, 13
atl_turning_angle, 14
atl_within_polygon, 15
